# *Botrytis cinerea* BcPTP1 is a late infection phase, cysteine rich protein cytotoxic effector

**DOI:** 10.1101/2021.07.18.452873

**Authors:** Wenjun Zhu, Mengxue Yu, Ran Xu, Kai Bi, Chao Xiong, Zhiguo Liu, Amir Sharon, Daohong Jiang, Mingde Wu, Qiongnan Gu, Ling Gong, Weidong Chen, Wei Wei

## Abstract

*Botrytis cinerea* is a broad-host-range necrotrophic phytopathogen responsible for serious crops diseases. To facilitate infection, *B. cinerea* secretes a large number of effectors that induce plant cell death. In screening secretome data of *B. cinerea* during infection stage, we identified a phytotoxic protein (BcPTP1) that can also induce immune resistance in plants. BcPTP1 is a small (90 aa), cysteine rich protein without any known domains. Transiently expression of BcPTP1 in leaves caused chlorosis that intensifies with time and eventually lead to cell death. Point mutations in eight of the 10 cysteine residues of BcPTP1 abolished the toxic effect, however residual toxic activity remained after heating the peptide, suggesting contribution of unknown epitopes to protein phytotoxic effect. The transcript level of the *bcptp1* gene was low during the first 36 h after inoculation and increased sharply upon transition to the late infection stage, suggesting a role of BcPTP1 in lesion spreading. While statistically insignificant, deletion of the *bcptp1* gene led to slightly smaller lesions on bean leaves. Further analyses indicated that BcPTP1 is internalized into plant cells after secreting into the apoplast and its phytotoxic effect is negatively regulated by the receptor-like kinases BAK1 and SOBIR1. Collectively, our findings show that BcPTP1 is a virulence factor that toxifies the host cells and facilitates lesion spreading during the late infection stage.

## Introduction

During the evolutionary arms race with phytopathogens, plants have developed a sophisticated defense system (Jones and Dangl, 2006). The first defense layer is pathogen-associated molecular patterns (PAMPs)-triggered immunity (PTI), in which host receptors recognize molecules or domains that are conserved in certain groups of pathogens and induce an effective resistance response (Albert *et al.,* 2020). In fungi and oomycetes, the best studied PAMPs include glucans (Fliegmann *et al.,* 2004), xylanase EIX (Rotblat *et al.,* 2002), chitin (Shinya *et al.,* 2015), endopolygalacturonases (Zhang *et al.,* 2014b), Pep13 (Brunner *et al.,* 2002), Cerato-platanin proteins (Yang *et al.,* 2018) and INF1 (Kanneganti *et al.,* 2006). In addition, several PAMPs that induce plant necrosis or PTI across different classes of microbes have also been well characterized, such as glycoside hydrolase 12 proteins (GH12) in the fungal pathogen *Verticillium dahliae* and *Botrytis cinerea* (Gui *et al.,* 2017; Zhu *et al.,* 2017) and the oomycete pathogen *Phytophthora sojae* (Ma *et al.,* 2015; Ma *et al.,* 2017; Wang *et al.,* 2018), Ave1 protein in multiple plant pathogenic fungi and bacteria (Thomma *et al.,* 2011; de Jonge *et al.,* 2012), and the classical necrosis and ethylene-inducing peptide 1-like proteins (NLPs) in multiple prokaryotic and eukaryotic microbial pathogens (Oome *et al.,* 2014; Albert *et al.,* 2015).

To overcome the basal plant immunity and infect hosts effectively, biotrophic and hemibiotrophic pathogens secrete diverse effectors to the interface area between fungal hyphae and host or into plant cells (Stergiopoulos and de Wit, 2009; Lo Presti *et al.,* 2015; Kim *et al.,* 2016). To cope with pathogen effectors, plants have co-evolved a second line of defense, also called effector-triggered immunity (ETI), which is mediated by resistance (R) proteins and is often associated with the local plant cells hypersensitive response (HR) at the infection site (Cui *et al.,* 2015). HR is efficient in restricting biotrophic and hemibiotrophic pathogens (Jones and Dangl, 2006; van Ooijen *et al.,* 2007; Cui *et al.,* 2015), but is ineffective against necrotrophic pathogens, which can turn it against the host to facilitate infection (Lorang *et al.,* 2012; Gao *et al.,* 2015).

Compared to the large number and well characterized effectors in biotrophic and hemibiotrophic fungal pathogens, relatively fewer effectors have been studied in necrotrophic fungal pathogens, especially the broad-host-range necrotrophic fungal pathogens. *Botrytis cinerea* is a typical broad-host-range necrotrophic fungal phytopathogen, causing gray mold and rot diseases in hundreds of plant species including many agriculturally important crops, and leading to enormous economic losses each year (Dean *et al.,* 2012; Veloso and van Kan, 2018). It was shown that the infection process of *B. cinerea* on host plant includes three typical stages: an early stage characterized by local necrosis lesions formation, an intermediate stage during which a variety of sophisticated interactions between plant-pathogen occur and determine the outcome of *B. cinerea* infections, and the late stage of fast-spreading lesions (Eizner *et al.,* 2017). Since *B. cinerea* infects and colonizes by killing plant cells, it secretes diverse effector proteins to manipulate the host defenses and/or to induce death for facilitating infection on host plants (Heard *et al.,* 2015; Frías *et al.,* 2016; Zhu *et al.,* 2017; Denton-Giles *et al.,* 2020; Frías *et al.,* 2011; Shao *et al.,* 2021). These secreted effector proteins play important roles during *Botrytis*-plant interactions. However, compared with biotrophic and hemibiotrophic pathogens effectors, the underlying biochemical activities and molecular mechanisms of the necrotrophic secreted proteins remain incompletely understood.

Aiming to characterize potential effectors of *B. cinerea*, we identified a secreted protein with a phytotoxic activity that was named BcPTP1 (<u>p</u>hytotoxic <u>p</u>rotein). Similar to most other cell death inducing proteins (CDIPs, Li *et al.,* 2020; Shao *et al.,* 2021), BcPTP1 functions in apoplastic space of plant cell and induces a plant defense response. However, unlike the vast majority of CDIPs, BcPTP1 leads to development of gradual chlorosis rather than instant cell death and the *bcptp1* gene is expressed late during infection and affects lesion development, namely it is a late-stage virulence factor.

## Results

### BcPTP1 is a secreted protein with phytotoxic activity

The effect of putative effector proteins was tested by *Agrobacterium tumefaciens*-mediated transient expression (agroinfiltration) of the candidate proteins fused with GFP tag at the C-terminus in *N. benthamiana* leaves. We found that the protein BcPTP1 (BCIN_05g03680)-GFP containing a potential N-terminal signal peptide caused chlorosis in *N. benthamiana* leaves that developed within 10 days after agroinfiltration, eventually leading to cell death within 15 days after agroinfiltration (Fig. 1). Transient expression of BcPTP1^ΔSP^-GFP without the secretion signal peptide did not trigger leaf chlorosis, indicating that BcPTP1 may function in the leaf apoplastic space. This phenotype differs from the induction of necrotic cell death that is common to previously characterized apoplastic CDIPs (Li *et al.,* 2020; Shao *et al.,* 2021), and suggests that BcPTP1 has a phytotoxic activity rather than a necrotic cell death inducing activity.

**Fig. 1.**
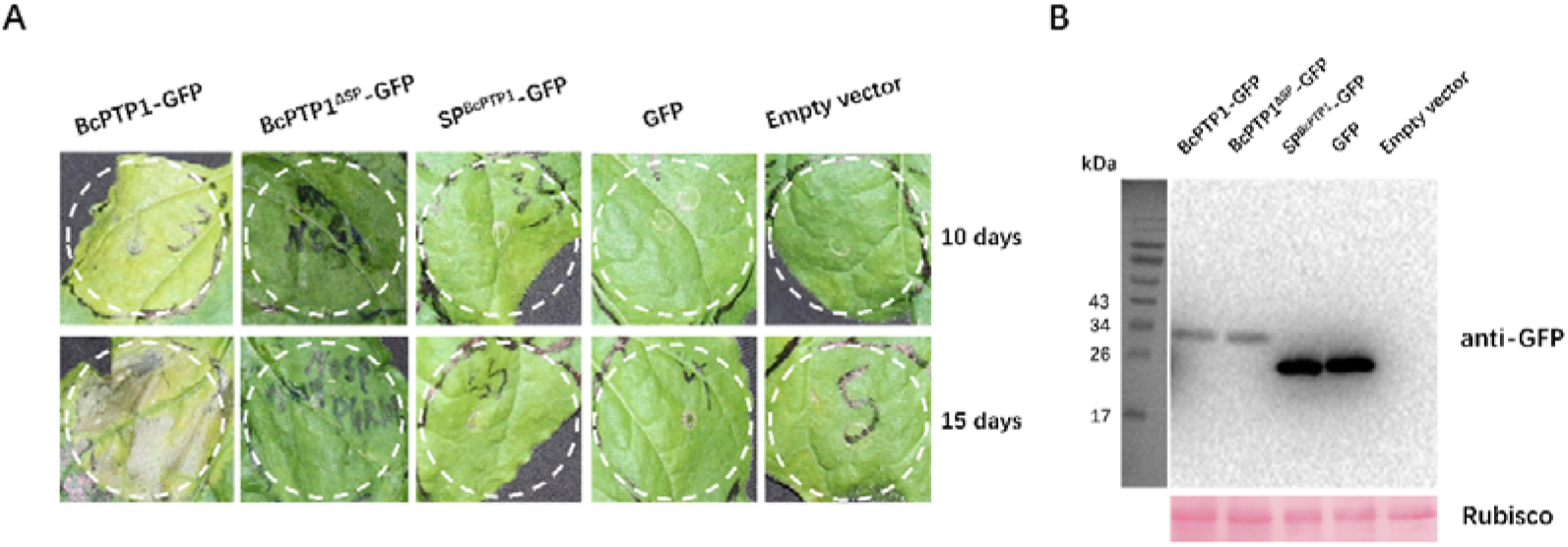
BcPTP1 is a phytotoxic secreted protein. (A) Images of *N. benthamiana* leaves 10 days and 15 days after agroinfiltration with *A. tumefaciens* strains carrying the indicated constructs. The *A. tumefaciens* carrying the empty vector was used as control. BcPTP1-GFP, BcPTP1 fused with GFP at the C-terminus; BcPTP1^ΔSP^-GFP, BcPTP1 without signal peptide fused with GFP at the C-terminus; SP^BcPTP1^-GFP, GFP fused with the signal peptide of BcPTP1. (B) Immunoblot analysis of proteins from *N. benthamiana* leaves transiently expressing the indicated constructs. Top panel, immunoblot using anti-GFP antibody; bottom panel, staining of the Rubisco large subunit with Ponceau S.

Two methods were used to verify whether BcPTP1 is a secreted protein. First, the BcPTP1 signal peptide (initial N terminus 19 amino acids) was used to replace the N-terminal signal peptide of BcXYG1, the strong death-inducing secreted protein of *B. cinerea* which functions in the apoplastic space (Zhu *et al.,* 2017), to produce SP^(BcPTP1)^-BcXYG1^ΔSP^-HA. The BcXYG1-HA with its native signal peptide was used as a positive control, and the BcXYG1^ΔSP^-HA, GFP-HA, SP^(BcPTP1)^ -GFP-HA and empty vector were used as negative controls. Within five days after agroinfiltration, ^ΔSP^ both of BcXYG1-HA and SP^(BcPTP1)^-BcXYG1-HA fusion proteins, which are secreted from the plant cells to the extracellular space, triggered typical plant cell death, whereas BcXYG1^ΔSP^ -HA, which lacks the signal peptide and therefore remains inside the plant cell, failed to induce cell death (Fig. 2A). In the second method, we fused the signal peptide of BcPTP1 to GFP to form SP^(BcPTP1)^-GFP. Likewise, we fused the signal peptide of BcXYG1 to GFP to form SP^(BcXYG1)^-GFP for comparison. The constructs were used to transform the wild type *B. cinerea* strain to generate overexpression strains of these fusion proteins. All the indicated proteins were successfully expressed in the hyphae of each strain as shown in western blot analysis (Fig. 2C). Then, all examined strains were cultured in potato dextrose broth (PDB) for 3 days, the culture filtrates were collected, purified and presence of the fused proteins was checked by western blot analysis. The results confirmed the accumulation of GFP protein in the culture medium of SP^(BcPTP1)^-GFP and SP^(BcXYG1)^-GFP overexpression strains, but not the GFP overexpression or wild type (WT) strain (Fig. 2D). Collectively, these analyses confirmed the secretion function of the BcPTP1 signal peptide.

**Fig. 2.**
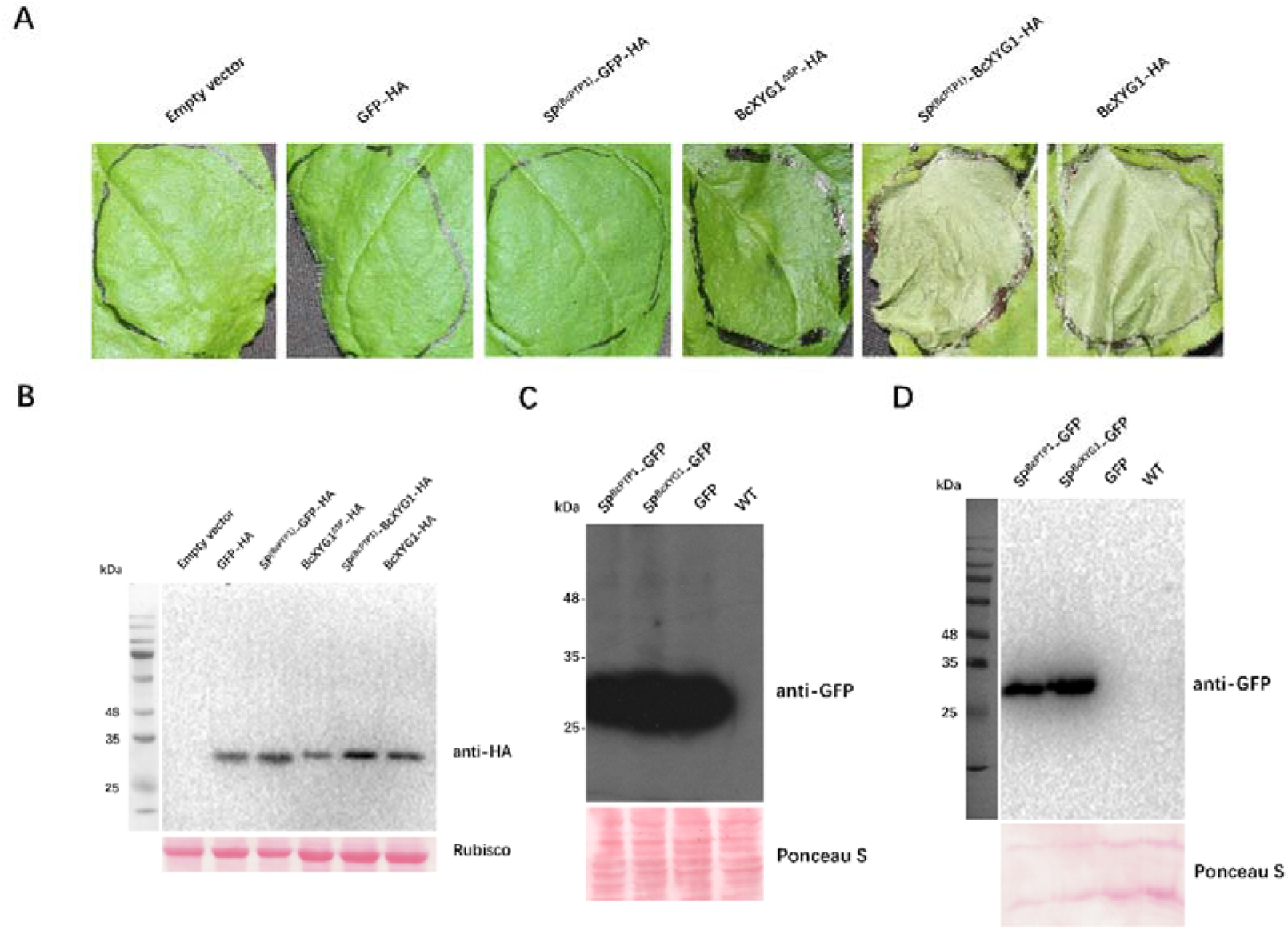
BcPTP1 signal peptide possesses secretion function. (A) Images of *N. benthamiana* leaves 5 days after agroinfiltration with *A. tumefaciens* strains carrying the indicated constructs. The *A. tumefaciens* carrying the empty vector was used as control. GFP-HA, HA-tagged GFP; SP^(BcPTP1)^-GFP-HA, HA-tagged GFP with the signal peptide of BcPTP1; BcXYG1^ΔSP^-HA, HA-tagged BcXYG1 lacking the native signal peptide; SP^(BcPTP1)^-BcXYG1-HA, HA-tagged BcXYG1 with its native signal peptide replaced with the BcPTP1 signal peptide; BcXYG1-HA, HA-tagged BcXYG1 with its native signal peptide. (B) Immunoblot analysis of proteins from *N. benthamiana* leaves transiently expressing the indicated constructs. Top panel, immunoblot using anti-HA antibody; bottom panel, staining of the Rubisco large subunit with Ponceau S. (C-D) The indicated genes were expressed under the regulation of *B. cinerea* histone H2B promoter (NCBI identifier CP009806.1) and the endo-β-1,4-glucanase precursor terminator (NCBI identifier CP009807.1). The *B. cinerea* strains were cultured in PDB medium for 72 h, then the culture filtrate was collected and purified by filtration. (C) Immunoblot analysis of total mycelia proteins from indicated *B. cinerea* strains. Top panel, immunoblot using anti-GFP antibody; bottom panel, Ponceau S staining of the total mycelia proteins. (D) Immunoblot analysis of culture filtrate proteins from indicated *B. cinerea* strains. Top panel, immunoblot using anti-GFP antibody; bottom panel, Ponceau S staining of the secretory proteins. SP^BcPTP1^-GFP, GFP fused with the signal peptide of BcPTP1; SP^BcXYG1^-GFP, GFP fused with the signal peptide of BcXYG1; GFP, GFP overexpression strain; WT, wild type strain.

### BcPTP1 is internalized into plant cell

The above results showed that BcPTP1 is a secreted protein and its phytotoxic activity depends on localization in the leaf apoplastic space (Fig. 1; Fig. 2). Thus, we postulated that BcPTP1 remains the in apoplastic space after secretion by the fungus. To verify this hypothesis, we analyzed the subcellular localization of BcPTP1-GFP fusion protein following agroinfiltration. Unexpectedly, we found that the BcPTP1-GFP fusion protein was mainly localized at the cytoplasmic vesicles and in the periphery of plant cell plasma membrane, whereas the BcPTP1^ΔSP^-GFP and GFP were mainly distributed in the nuclei and cytoplasm, and the SP^BcPTP1^-GFP was concentrated in the apoplastic space (Fig. 3). The unexpected localization of BcPTP1 indicates that besides phytotoxic activity in apoplast, BcPTP1 can be also internalized into plant cells for other unknown functions after the initial secretion to apoplastic space.

**Fig. 3.**
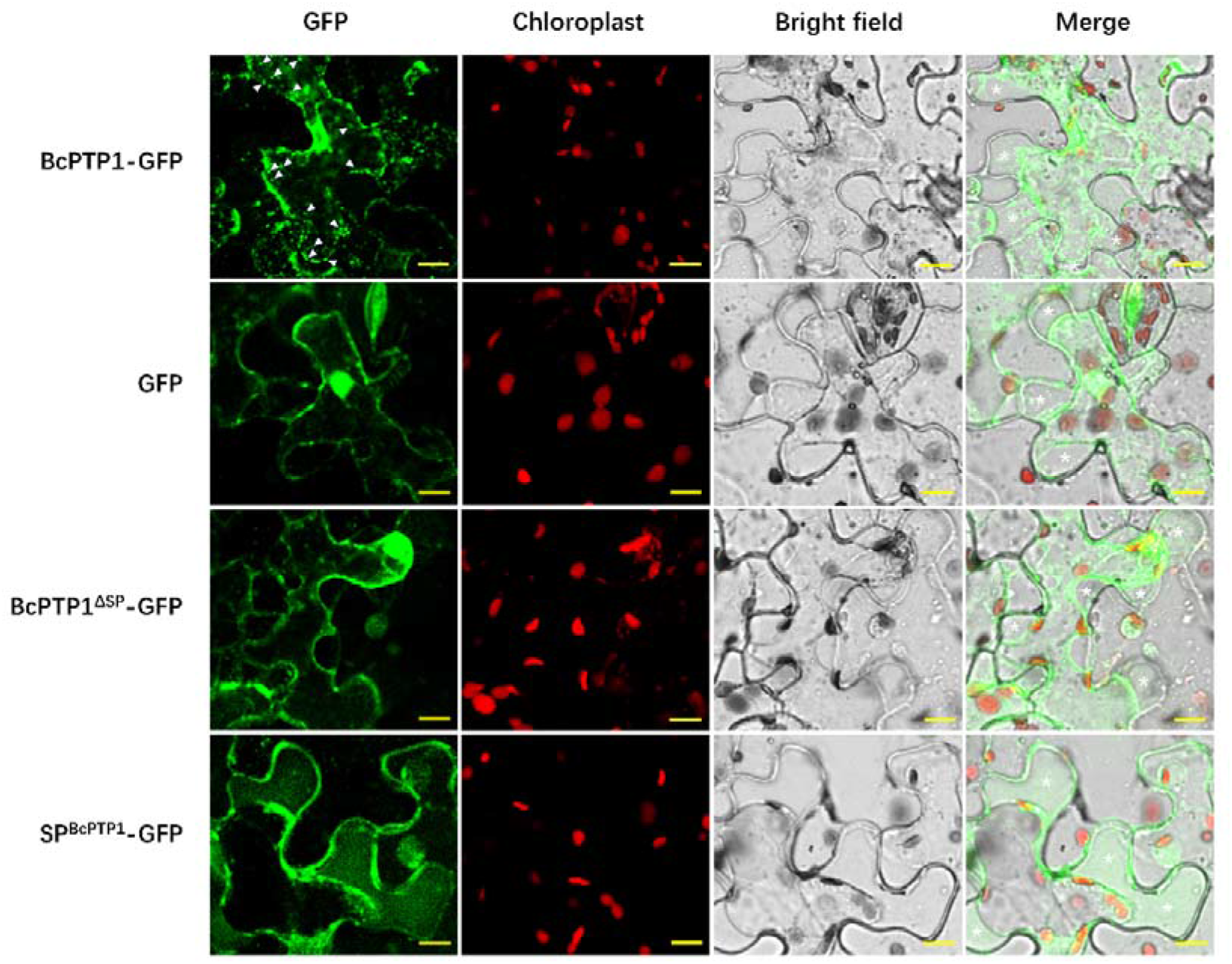
Subcellular localization of BcPTP1 in *N. benthamiana* epidermal cells. Leaves epidermal cells were harvested 3 days after agroinfiltration and plasmolyzed in 0.75 M sucrose solution, then the samples were scanned by confocal laser scanning microscope. BcPTP1-GFP, GFP fused to full length BcPTP1; BcPTP1^ΔSP^ -GFP, GFP fused to BcPTP1 without signal peptide; SP^BcPTP1^-GFP, GFP fused to the BcPTP1 signal peptide. The white arrows indicate the internalized cytoplasmic vesicle. White asterisks mark apoplastic space of plant cells. Bars = 10 μm

### BcPTP1 is toxic to dicot but not monocot plants

To determine whether BcPTP1 affects plants other than *N. benthamiana* and to avoid the *A. tumefaciens* incompatibility issues on plants, we produced and purified the BcPTP1 protein from *Escherichia coli* (Supplementary Fig. S1). Infiltration assay with different concentrations of the purified protein demonstrated that 25 μg/ml of BcPTP1 recombinant protein was sufficient to trigger leaf chlorosis in *N. benthamiana* (Fig. 4A). Cell death developed following treatment of the leaves with higher protein titers (50-100 μg/ml). In addition, BcPTP1 also induced cell death in tomato and *A. thaliana* leaves, but not in the monocot maize even at high protein concentrations (Fig. 4B). These results indicated that BcPTP1 is toxic to multiple dicot plants, but not to monocot cereal.

**Fig. 4.**
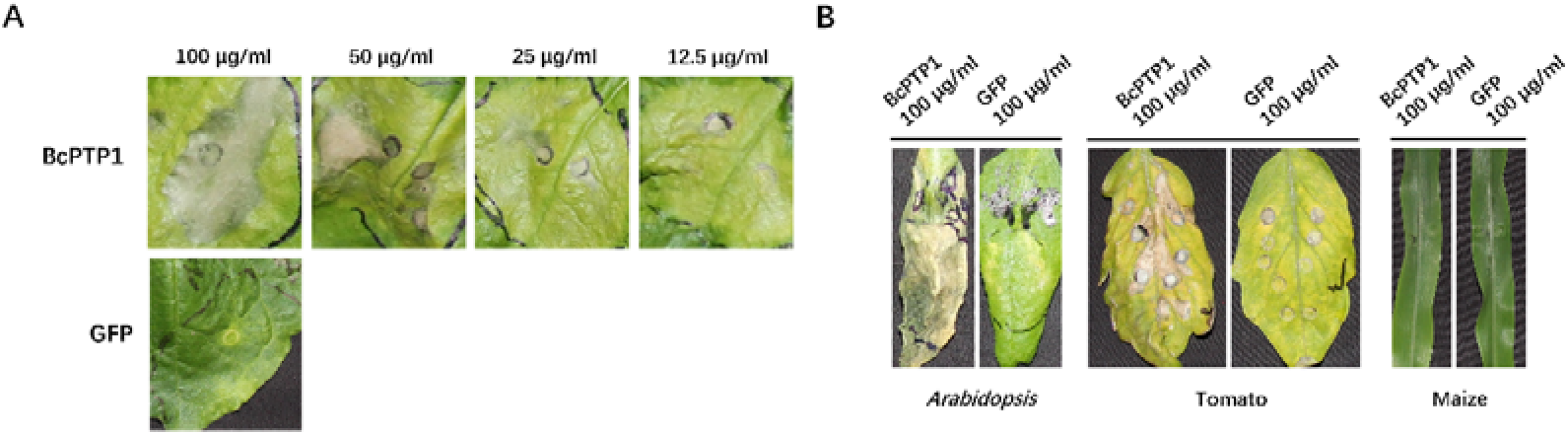
The BcPTP1 protein is phytotoxic to dicot but not monocot plant species. Proteins were produced in *E. coli*, purified and suspended in phosphate-buffered saline (PBS). (A) Response of *N. benthamiana* leaves infiltrated with different concentrations of BcPTP1 at 15 days after treatment. (B) Response of Arabidopsis, tomato and maize leaves infiltrated with 100 μg/ml protein solution.

### Cysteine residues in BcPTP1 are important for protein toxicity

*bcptp1* is a single-copy gene in *B. cinerea*, encoding 90 amino acids with 10 cysteine residues, out of which eight residues are highly conserved (Supplementary Fig. S2). Bioinformatics analysis demonstrated that the first 20 N-terminal amino acids encode a signal peptide. No other known protein domains or possible functions were predicted using ***SMART MODE*** analysis (http://smart.embl-heidelberg.de/smart/change_mode.pl), and also no nuclear localization signal or chloroplast transit peptide were found in protein sequence. BLAST searches against the **NCBI** database with the BcPTP1 sequence as query showed that homologs of BcPTP1 with high similarity are only present in a small number of fungal genera, most of which are plant pathogens, including *Botryotinia*, *Sclerotinia*, *Alternaria*, *Bipolaris*, *Fusarium*, *Colletotrichum*, *Botrytis,* and saprotrophic *Aspergillus* species. Significantly, no homologs were found in any of the biotrophic plant pathogens or in human pathogenic fungi. Multiple sequence alignment and phylogenetic analysis of BcPTP1 and its homologues showed significant sequence similarity (Supplementary Fig. S2A and S2B). 3D structure prediction of BcPTP1 using *I-TASSER* showed that BcPTP1 contains two α-helices on each side of the protein structure and two internal β-strands (Supplementary Fig. S2C).

Many effectors are small secreted cysteine-rich proteins (SSCP). The cysteine residues contribute to the formation of disulfide bonds (Sevier and Kaiser, 2002; Marianayagam *et al.,* 2004), which are essential for the structure and function of these proteins (Stergiopoulos and de Wit, 2009). To examine whether phytotoxicity of BcPTP1 depends on its cysteine residues, we replaced each of the 10 cysteine residues with alanine individually using site-directed mutagenesis. *A. tumefaciens* infiltration assay of *N. benthamiana* leaves demonstrated that mutations of C26A or C34A individually could induce more severe phenotype than the native BcPTP1 protein. The activity of the C67A mutant decreased compared to the native protein, whereas mutations in any of the seven remaining cysteine residues completely abolished phytotoxicity (Fig. 5). Additionally, boiling the BcPTP1 protein at 100□ for 30 min reduced, but did not completely abolished phytotoxicity (Fig. 6). These results suggest that certain epitopes, which are affected by specific cysteine residues, mediate the phytotoxic activity of the BcPTP1 protein.

**Fig. 5.**
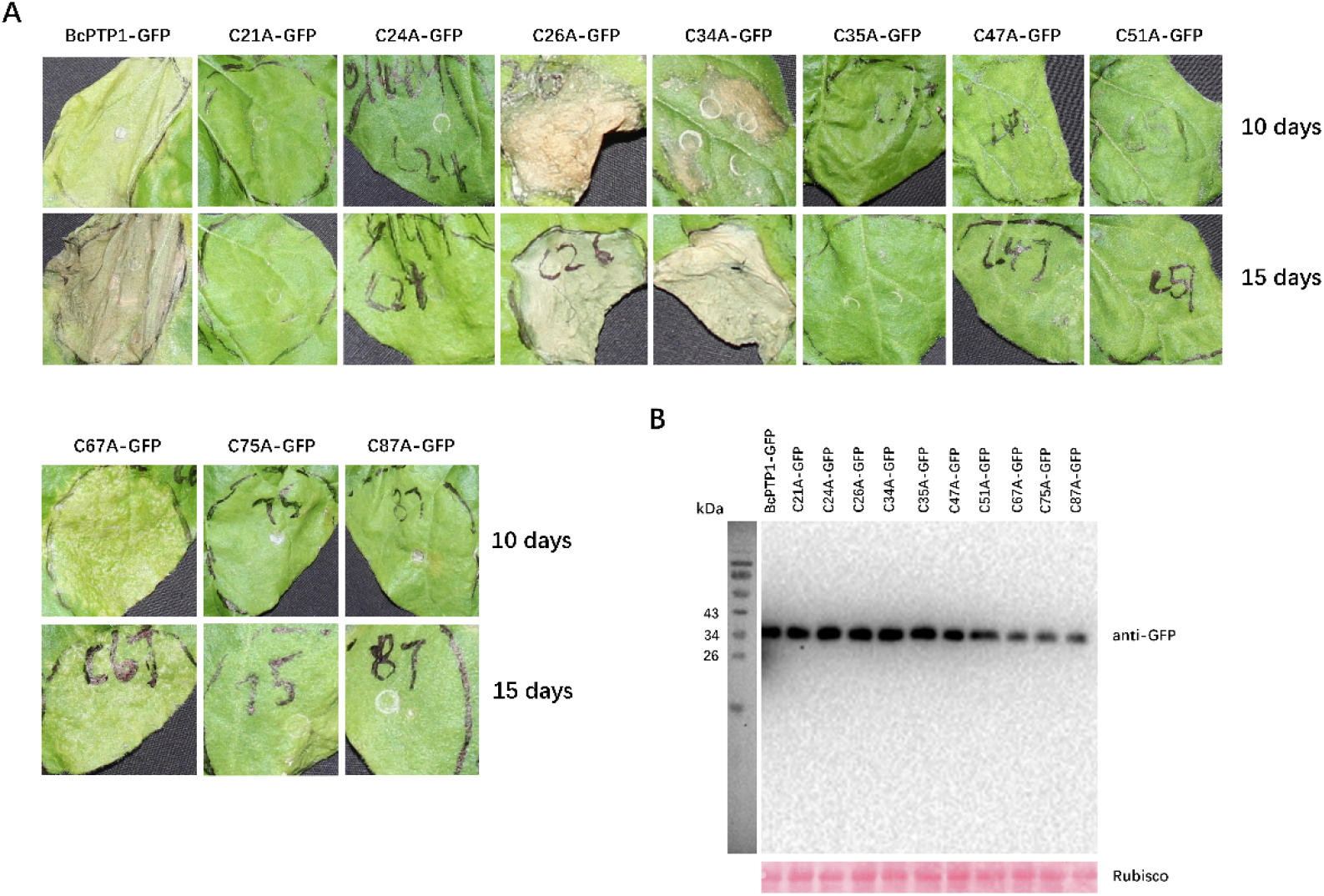
Specific cysteine residues necessary for BcPTP1 phytotoxicity. (A) Images of *N. benthamiana* leaves 10 days and 15 days after transient agroinfiltration. (B) Immunoblot analysis of proteins from *N. benthamiana* leaves transiently expressing the cysteine residue mutant constructs. Top panel, immunoblot using anti-GFP antibody; Bottom panel, staining of the Rubisco large subunit with Ponceau S.

**Fig. 6.**
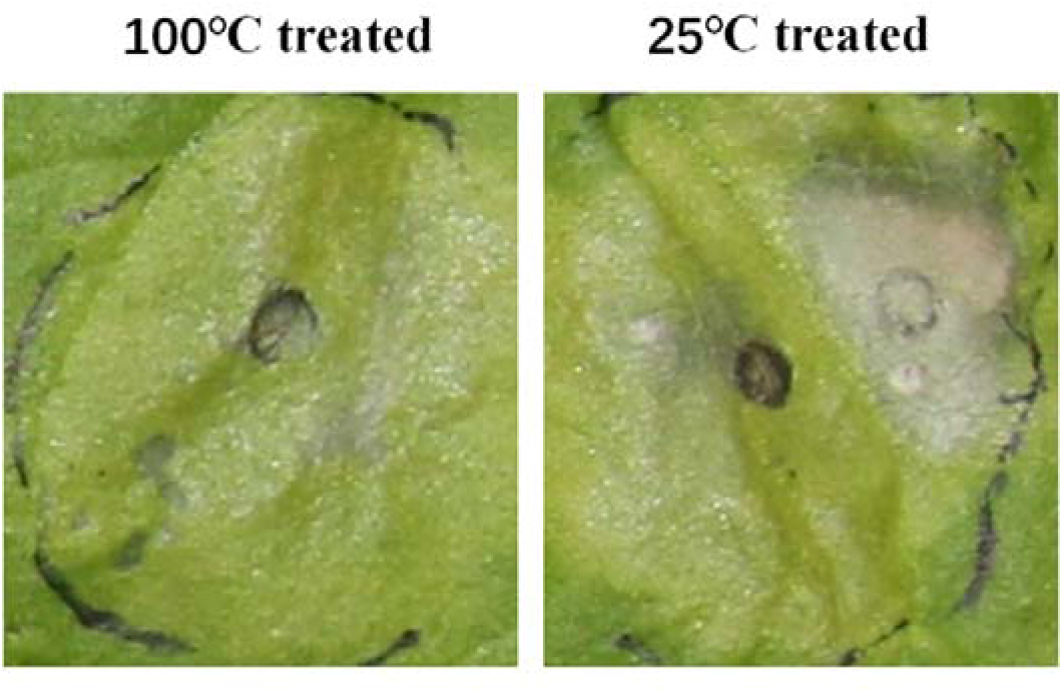
Phytotoxicity of BcPTP1 is partially heat-stable. Images of *N. benthamiana* leaves 15 days after infiltration with 50 μg/ml of heat-treated (100□ treated for 30 min) and native (25□ treated for 30 min) BcPTP1 protein.

### *bcptp1* is highly expressed at late infection stage but is not essential for pathogenicity

To analyze the biological roles of BcPTP1 during *B. cinerea* infection, we measured the expression levels of the *bcptp1* gene during infection. The transcript levels of *bcptp1* were low at early infection stage and then increased sharply about 48 hpi with a peak at 60 hpi, about 35-fold higher than at 0 hpi (Fig. 7). When *B. cinerea* was cultured on solid Gamborg’s B5 medium, the transcript level of *bcptp1* was increased about 12-fold at 36 hpi compared to earlier time points, and then remained stable throughout the culture period (Fig. 7).

**Fig. 7.**
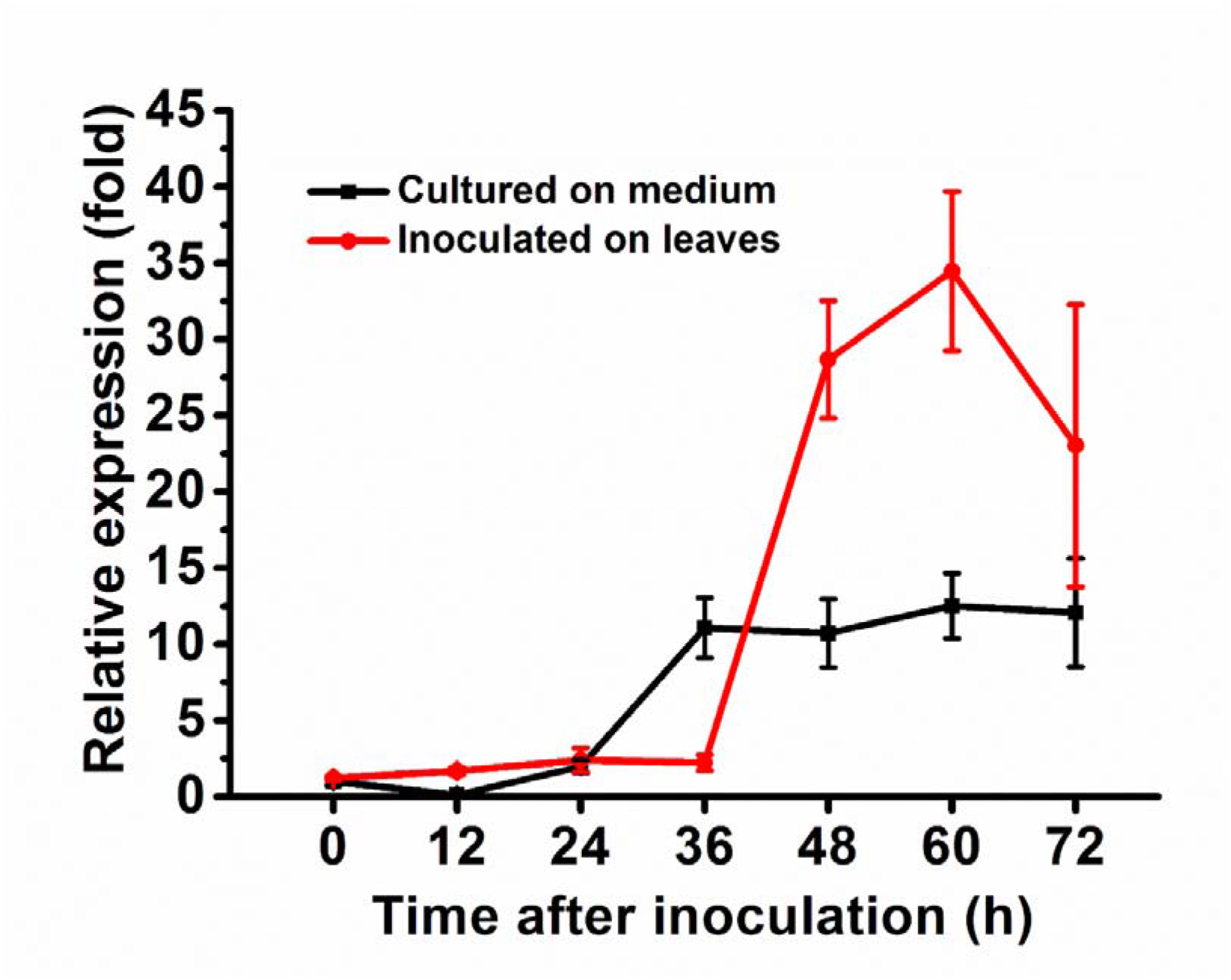
Expression of the *BcPTP1* gene is up-regulated at late infection stage. Bean leaves (red line) or Gamborg’s B5 medium (black line) were inoculated with *B. cinerea* conidia, and expression levels of the *BcPTP1* gene were evaluated by RT-qPCR. The expression level of *BcPTP1* inoculated on plant or in Gamborg’s B5 medium at 0 h was set as 1, and relative transcript levels were calculated using the comparative Ct method. Transcript levels of the *B. cinerea Bcgpdh* gene were used to normalize different samples. Data represent means and standard deviations of three independent replications.

To further investigate possible role of BcPTP1 in *B. cinerea* pathogenicity, we generated *bcptp1* gene deletion and overexpression strains (Supplementary Fig. S3). All the deletion and overexpression strains had normal colony morphology, conidia production and growth rate on potato dextrose agar (PDA) (Supplementary Fig. S4). In addition, we did not observe obvious changes in sensitivity of the transgenic strains to various types of stresses, including 1 M NaCl, 1 M sorbitol, 0.02% SDS, 20 mM H_2_O_2_, 0.3 mg/ml Calcofluor White and 0.5 mg/ml Congo Red (Supplementary Fig. S5).

Pathogenicity study showed that the *bcptp1* deletion strains caused slightly smaller disease lesion on bean leaves than the wild type strain, although the differences were statistically insignificant (Supplementary Fig. S6). Likewise, the *bcptp1* over expression strains did not show obvious difference in lesion size (Supplementary Fig. S6). The results indicated that deletion or overexpression of *bcptp1* does not significantly affect the final outcome of *B. cinerea* infection.

### BcPTP1 triggers immune response and induces resistance against *B. cinerea* in *N. benthamiana*

High proportion of the analyzed CDIPs are recognized by the plant immune system and activate a defense response. To analyze whether BcPTP1 can induce plant resistance to *B. cinerea*, *N. benthamiana* leaves were infiltrated with 10 μg/ml of purified BcPTP1 or GFP proteins, 48 h later, the infiltrated leaves were inoculated with *B. cinerea* mycelia plugs, the plants were incubated for an additional 48 h in a moist chamber and then symptoms were recorded. The results showed that the infection on tobacco leaves pre-treated with BcPTP1 protein was significantly reduced compared to leaves pre-treated with GFP protein (Fig. 8A), indicating that along with phytotoxicity, BcPTP1 can also trigger plant resistance.

**Fig. 8.**
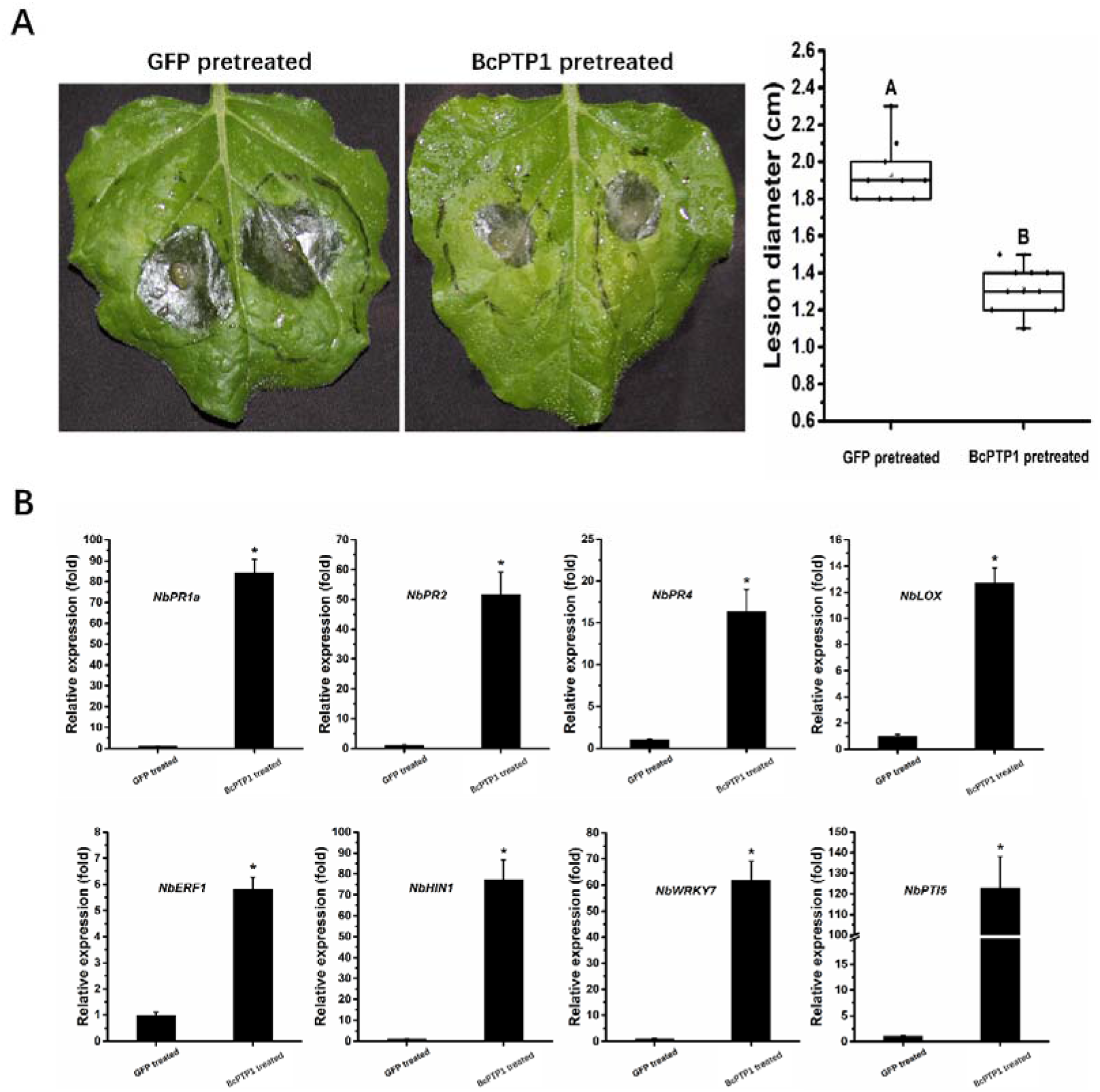
BcPTP1 induces resistance in tobacco. (A) *N. benthamiana* leaves were infiltrated with 10 μg/ml purified BcPTP1 or GFP protein. After 2 days, the infiltrated leaves were inoculated with *B. cinerea* in a humid chamber. The lesions were photographed and measured at 48 hpi. Data were obtained from three independent experiments with total 10 samples. In box plots, whiskers indicate the minimum and maximum values, the line indicates the median, the box boundaries indicate the upper (25th percentile) and lower (75th percentile) quartiles, all data are plotted as black dots. Different letters in the graph indicate statistical differences at P ≤ 0.01 using ANOVA (one-way). (B) Relative expression levels of defense-related genes from *tobacco* leaves that treated with BcPTP1 or GFP for 48 h were determined by RT-qPCR analysis. The expression level of indicated genes in GFP-treated leaves were set as 1. The expression level of the tobacco *NbEF1α* gene was used to normalize different samples. Data represent means and standard deviations of three independent replicates. Asterisks in the graph indicate statistical differences at P ≤ 0.01 using ANOVA (one-way).

To verify whether the enhanced resistance of tobacco caused by BcPTP1 is associated with the expression changes of defense-related genes, we analyzed the transcript alteration of salicylic acid (SA) signal pathway genes *NbPR1a* and *NbPR2*, Jasmonic acid (JA) signal pathway genes *NbPR4* and *NbLOX*, ethylene signal pathway gene *NbERF1*, HR-related genes *HIN1*, and PTI-related genes *NbWRKY7* and *NbPTI5* using RT-qPCR, as previously described (Nie *et al.,* 2019). In all cases we observed a drastic increase in gene expression after infiltration of the leaves with 10 μg/ml BcPTP1 (Fig. 8B).

### BAK1 and SOBIR1 negatively regulate the death-inducing activity of BcPTP1 in *N. benthamiana*

Since BcPTP1 is targeted to the apoplast space of *N. benthamiana* tissue, it is possible that similar to other CDIPs, BcPTP1 interacts with plant membrane receptor-like proteins (RLPs) to transmit the immunity signals via the RLP-SOBIR1-BAK1 complex (Liebrand *et al.,* 2013; Zhang *et al.,* 2014b; Albert *et al.,* 2015; Ma *et al.,* 2015; Postma *et al.,* 2016; Gui *et al.,* 2017; Zhu *et al.,* 2017). To test this possibility, we generated *NbBAK1*- and *NbSOBIR1*-silenced *N. benthamiana* plants using virus-induced gene silencing (VIGS). The gene-silenced plants were agroinfiltrated with BcPTP1 expression construct. Unexpectedly, BcPTP1 induced a much more severe phytotoxicity in the *BAK1*- or *SOBIR1*-silenced plants than in wild type tobacco plants (Fig. 9). The result indicated that the death-inducing signal of BcPTP1 is mediated through an unknown signal transduction pathway, and is negatively regulated by BAK1 and SOBIR1.

**Fig. 9.**
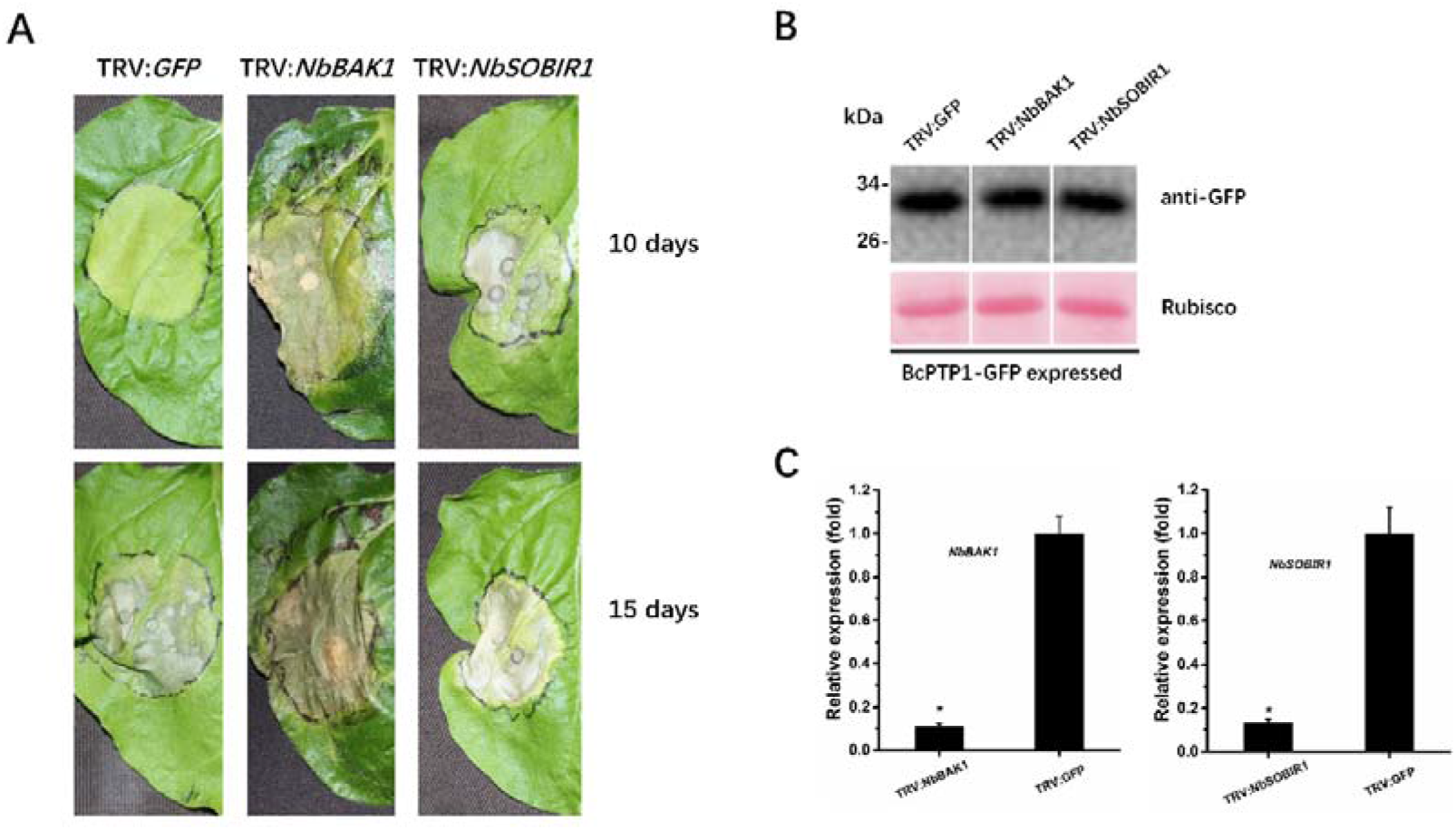
BAK1 and SOBIR1 negatively regulate the death-inducing activity of BcPTP1 in *N. benthamiana.* TRV-based VIGS vectors were used to initiate silencing of *NbBAK1* (TRV : *NbBAK1*) and *NbSOBIR1* (TRV : *NbSOBIR1*). TRV : *GFP* was used as a control virus treatment in these experiments. (A) Three weeks after initiation of VIGS, *BcPTP1-GFP* was transiently expressed in the gene-silenced leaves using agroinfiltration. Leaves were photographed 10 and 15 days after treatment. (B) Immunoblot analysis of proteins from indicated *N. benthamiana* leaves transiently expressing BcPTP1-GFP. Top panel: BcPTP1-GFP was detected using anti-GFP antibody; bottom panel, staining of the Rubisco large subunit with Ponceau S. (C) *NbBAK1* and *NbSOBIR1* expression levels in gene-silenced tobacco leaves were determined by RT-qPCR analysis. Expression level in control plants (TRV : *GFP*) was set as 1. *NbEF1α* was used as an endogenous control. Data represent means and standard deviations from three biological replicates. Asterisks indicate significant differences (P ≤ 0.01).

## Discussion

The search for *B. cinerea* cell death-inducing effectors has yielded an increasing number of candidates DCIPs, most of which are associated with production of local necrosis during the early infection stage (Li *et al.,* 2020; Shao *et al.,* 2021). In search of factors that facilitate disease progression in late infection stage (lesion spreading), we identified BcPTP1, a SSCP with phytotoxic activity, which is expressed at the late infection stage. As such, BcPTP1 fulfills the definition of a late-stage necrotrophic effector.

The late *in planta* induction of *bcptp1* (Fig. 7) supports a potential role of BcPTP1 in the pathogenicity of *B. cinerea* at the late infection stage, probably as a factor that facilitates lesion expansion. Since the BcPTP1 is recognized by the plant immune system (Fig. 8), the late expression of the *bcptp1* gene might also prevent early perception and induction of a plant defense response. Despite the phytoxicity and induction *in planta*, deletion of *bcptp1* gene had only minor effect on fungal virulence (Supplementary Fig. S6). Such minor, or even lack of a visible change in virulence is common to many fungal effectors, and probably reflects presence of similar effectors with a redundant function or a quantitative effect that adds to the overall virulence arsenal of the fungus.

Compared to other well studied CDIPs, which possess strong activity and trigger death of tobacco leaves within less than 5 days (Ma *et al.,* 2015; Zhang *et al.,* 2017; Zhu *et al.,* 2017; Yang et al., 2018; Bi *et al.,* 2021; Yang *et al.,* 2021; Yin *et al.,* 2021), treatment of leaves with purified BcPTP1 protein or transient expression of using agroinfiltration, both induced chlorosis rather than instant cell death (Fig. 1; Fig. 4A). Hence, the BcPTP1 protein probably does not cause direct damage to the plant cell but more likely, it affects some unknown pathways that eventually lead to cell death, such as accumulation of toxic metabolites or inhibition of photosynthesis.

Homologues of BcPTP1 were found in several additional fungal species, mostly necrotrophic and hemibiotrophic plant pathogens (Supplementary Fig. S2). Significantly, no homologues are found in biotrophic plant pathogens or in human pathogens, which hint to a specific role of the protein in promoting necrotrophic infection. Homologues were also found in saprophytic *Aspergillus* species that colonize and grow on dead plant residues. While these saprotrophic fungi probably do not need the death-inducing activity of the BcPTP1 homologs during their entire life cycle, it is possible that they benefit from an unknown activity of these proteins, or they might have a yet unidentified plant associated phase.

Many effectors contain multiple cysteine residues, which form disulphide bonds (Sevier and Kaiser, 2002; Lu and Edwards, 2016). These residues play significant roles in protein folding, structural stability and protection of such proteins against degradation in harsh acidic and protease-rich environment when they are delivered into plant apoplast during infection (Rep, 2005). BcPTP1 contains 10 cysteine residues that are predicted to form potential disulphide bonds. Mutation of seven out of the 10 cysteine residues in BcPTP1 completely abolished, and an additional one greatly reduced, the phytotoxic activity, whereas mutation of two other residues enhanced the activity (Fig. 5A). It is possible that the majority of the cysteine residues are involved in forming the proper tertiary structure of BcPTP1, thus changing these cysteine residues abolished or decreased the death-inducing activity. The other two residues might be part of epitopes that contribute to the cell death-inducing activity and the mutations might lead to exposure of the immunogenic epitopes, resulting in enhanced activity. Similar study recently revealed that the secreted apoplastic protein PC2 from the potato late blight pathogen *Phytophthora infestans* was cleavage by plant apoplastic proteases, which is the essential process for PC2 to release the immunogenic peptides, thus to activate plant defense responses (Wang *et al.,* 2021a). Indeed, several studies demonstrated that heat denaturation or structural mutation of some secreted proteins did not abolished the necrosis-inducing activity (Zhang *et al.,* 2014a; Zhang *et al.,* 2014b), indicating that the potential epitopes are sufficient for the plant cell death-inducing activity. Similarly, our study also showed that heat-denatured BcPTP1 still retained a weak cytotoxic activity (Fig. 6). Accordingly, it is possible that in addition to tertiary structure, the unknown immunogenic peptide also contributes to the cell death inducing activity of BcPTP1.

Our study showed that BcPTP1 activates an immune response in *N. benthamiana* as measured by infection assay and induced expression of defense genes (Fig. 8). Thus, the BcPTP1 may also function as a potential elicitor. Indeed, our results demonstrated that BcPTP1 is secreted extracellularly (Fig. 1; Fig. 2) and may interact with plants receptor-like kinases (RLKs) and/or receptor like proteins (RLPs) and transmit signals via the RLKs BAK1 and/or SOBIR1, as other well-known elicitors (Albert *et al.,* 2015; Du *et al.,* 2015; Postma *et al.,* 2016; Franco-Orozco *et al.,* 2017; Zhu *et al.,* 2017; Nie *et al.,* 2019; Nie *et al.,* 2021). However, silencing of *BAK1* or *SOBIR1* in tobacco did not block cell death development but instead, it resulted in induction of more severe and faster plant cell death (Fig. 9). Similar studies had also been reported that some apoplast elicitors could induce necrosis independent of BAK1 and/or SOBIR1, such as BcPG3 (Zhang *et al.,* 2014b), TvEIX (Bar *et al.,* 2010), VdEIX3 (Yin *et al.,* 2021) and Fg12 (Yang *et al.,* 2021). It is possible that BcPTP1 and its putative interacting RLP may coordinate with other unknown LRR-RLKs as co-receptors to transmit immune or death-inducing signals.

Subcellular localization analysis of BcPTP1 showed that the fluorescence signal of BcPTP1-GFP fusion protein is mainly observed in plant cell cytoplasmic vesicles and periphery of plasma membrane, but not the expected apoplastic localization (Fig. 3). This intracellular localization of BcPTP1 is unexpected given that an extracellular localization of the protein is necessary for the cytotoxic activity. Several studies showed that the internalization of certain secreted fungal proteins was dependent on the plasma membrane RLKs BAK1 and/or SOBIR1 (Robatzek, 2006a, Robatzek *et al.,* 2006b; Chinchilla *et al.,* 2007a, Chinchilla *et al.,* 2007b; Liebrand *et al.,* 2013; Wang *et al.,* 2021b). Based on these studies and our findings, we proposed hypothetical model (Fig. 10) in which following secretion by the fungus, the BcPTP1 interacts in the apoplast with a plant RLK and/or RLP, and this complex transmits the phytotoxicity and defense signals. In parallel, the BcPTP1 protein might be also recognized by other plasma membrane receptor that cooperate with the BAK1 and SOBIR1 complex to mediate the internalization through cytoplasmic vesicles (Fig. 10). According to this model, silencing of the *BAK1* or *SOBIR1* genes blocks BcPTP1internalization leading to accumulation of large amounts of BcPTP1 in apoplastic space and increased phytotoxicity.

**Fig. 10.**
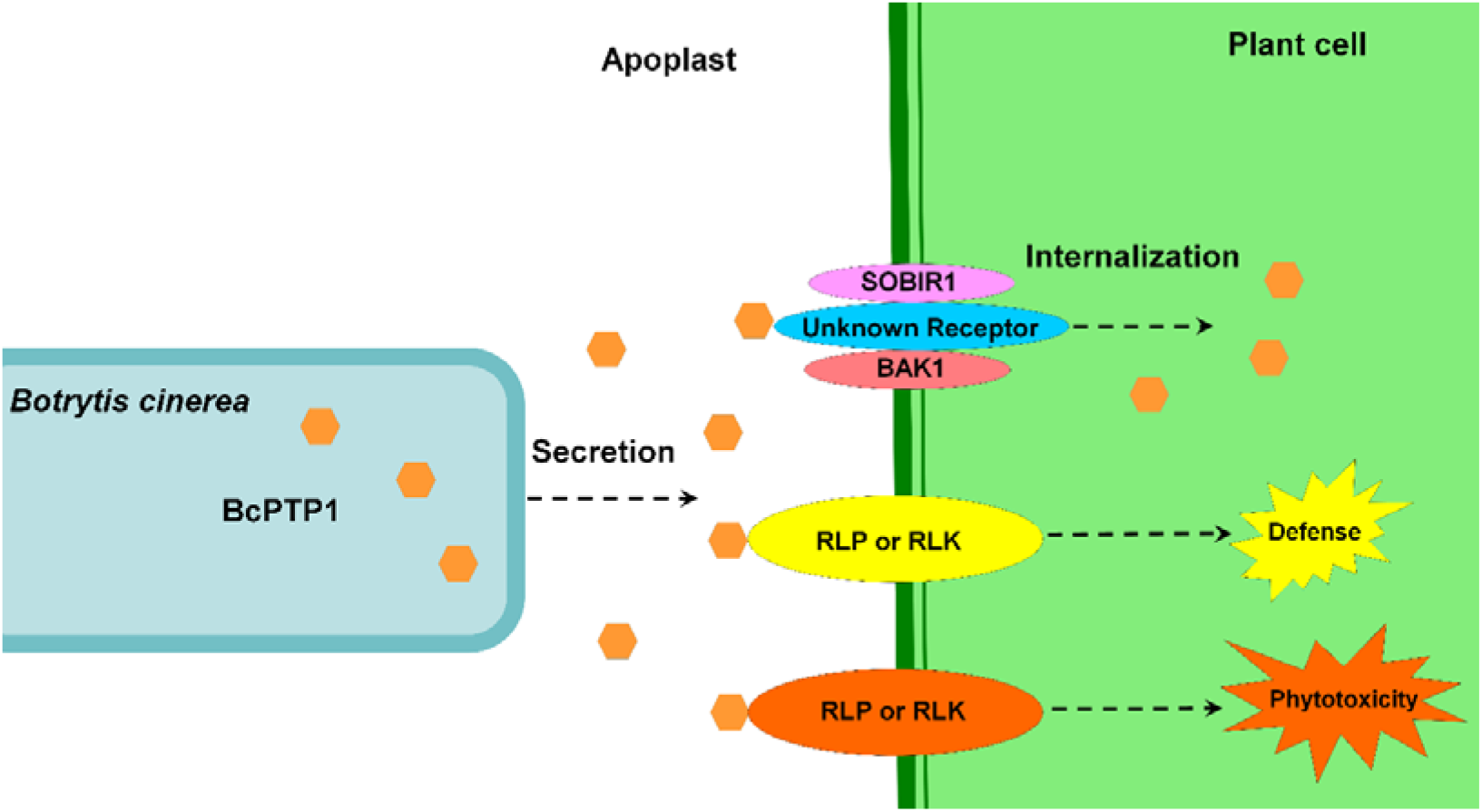
The model illustrating the internalization of BcPTP1 mediated by BAK1 and SOBIR1, and the defense and phytotoxicity induced by BcPTP1 though RLPs or PLKs in plant cell. BcPTP1 is secreted from *B. cinerea* hyphae into apoplastic space to interact with plants RLKs and/or RLPs and transmit the phytotoxicity and defense signals. BcPTP1 can also be recognized by other unknown receptor that cooperate with BAK1 and SOBIR1 complex to mediate the internalization.

## Acknowledgements

This research was supported by the National Natural Science Foundation of China (Grant No 31972215, 31861143043 and 81903782), Institute of Plant Protection and Soil Science, Hubei Academy of Agricultural Sciences/Key Laboratory of Integrated Pest Management on Crops in Central China, Ministry of Agriculture, P.R. China/Hubei Key Laboratory of Crop Diseases, Insect Pests and Weeds Control (Grant No 2018ZTSJJ14), and Hubei Province Agricultural Science and Technology Innovation Center Project (Grant No 2018-620-003-001).

## Author contributions

WZ and WW: conceptualization of experiments and research plans; WZ, MY, QG, LG and WW: performing the experiments; RX, KB, CX and ZL: data and statistical analysis; WZ, AS, DJ and MW: funding acquisition; AS and WC: manuscript revision; WZ and WW: writing.

## Data availability

Data of this study are all available within the paper and within its supplementary data published online. Further information may be obtained from the corresponding author.

## Materials and methods

### Fungi, bacteria, plants and culture conditions

The *Botrytis cinerea* wild type strain B05.10 and derived transformants used in this study were grown and maintained on PDA medium (Acumedia, MI, USA) at 22□ under continuous fluorescent light supplemented with near-UV (black) light. Conidia of all *B. cinerea* strains were obtained from 7-days-old cultures. *Escherichia coli* strains Rosetta-gami (DE3) and JM109 (Shanghai Weidi Biotechnology, Shanghai, China) were respectively used for proteins expression and plasmid construction. *Agrobacterium tumefaciens* strain GV3101(pSoup) (Shanghai Weidi Biotechnology, Shanghai, China) was used for *Agrobacterium*-mediated transient expression of target proteins in plant leaves.

Tobacco (*Nicotiana benthamiana*), bean (*Phaseolus vulgaris*, genotype N9059), maize (*Zea m*ays cv. Silver Queen) and tomato (*Solanum lycopersicum* cv. Hawaii 7981) plants were grown in a greenhouse under 16-h/8-h intervals of 25°C/22°C, light/dark. *Arabidopsis thaliana* Columbia-0 was grown in a chamber under 16-h/8-h intervals of 22°C/20°C, light/dark.

### Plasmid construction

The *bcptp1* gene deletion construct *Del-bcptp1* was generated as described previously (Zhu *et al.,* 2017) that the 5’ (510 bp) and 3’ (516 bp) flanks fragments of *bcptp1* gene were amplified and respectively cloned into the upstream and downstream regions of *hph* cassette using Gibson Assembly Master Mix kit (New England Biolabs, Massachusetts, USA). The overexpression plasmid OEBcPTP1- pH2G was generated that the full-length open reading frame of *bcptp1* was cloned into the pH2G vector under the regulation of *B. cinerea* histone H2B promoter (NCBI identifier CP009806.1) and the endo-β-1,4-glucanase precursor terminator (NCBI identifier CP009807.1), as described previously (Zhu *et al.,* 2017).

To transiently express the target protein in plant using agroinfiltration, the indicated sequences were cloned into binary plasmid pCNG between the 2×CaMV 35S promoter and NOS terminator (Yang *et al.,* 2018), then transformed into *A. tumefaciens* strain GV3101. The *E. coli* protein expression vectors were constructed that the GFP and *BcPTP1* mature sequence without the signal peptide were, respectively, cloned into vector pET-N-GST-PreScission (Beyotime Biotechnology, Shanghai, China), then transformed into *E. coli* strain Rosetta-gami (DE3). All the primers used for plasmid construction were listed in Table S1.

### Manipulation of nucleic acids

Total RNA of fungi and plant samples were isolated using RN03-RNApure Kit (Aidlab, Beijing, China), residual DNA was removed using DNase I (Thermo Scientific, MA, USA) according to manufacturers’ protocols and stored at –80°C. The first strand cDNA of indicated sample was generated using RevertAid First Strand cDNA Synthesis Kit (Thermo Scientific, MA, USA). The reverse transcription- quantitative PCR (RT-qPCR) was performed to analyze the gene expression profile using SYBR^®^ Green Supermixes (Bio-Rad, CA, USA) and CFX96 Touch Real-Time PCR Detection System (Bio-Rad, CA, USA) according to manufacturer’s instructions. The *B. cinerea Bcgpdh* gene and the *N. benthamiana NbEF1*α gene were, respectively, used as endogenous control genes for normalizing the expression levels of target genes as described previously (Zhu *et al.,* 2017). Primers were designed across or flanking an intron (Supplementary Table S1). For each analyzed gene, RT-qPCR assays were repeated at least twice, each repetition with three independent replicates. The genomic DNA of indicated *B. cinerea* strains were isolated using Fungal Genomic DNA Purification Kit (Simgen, Hangzhou, China) according to the manufacturer’s protocol.

### Bioinformatics analysis

The NCBI (http://www.ncbi.nlm.nih.gov/) database was used to obtain homologous sequences of BcPTP1 from other pathogens using BLASTp analysis. The JGI (http://genome.jgi.doe.gov/Botci1/Botci1.home.html) database of *B. cinerea* was used to characterize *B. cinerea* genomic and transcriptomic sequences. The HMMSCAN (https://www.ebi.ac.uk/Tools/hmmer/search/hmmscan) was used to analyze the protein domain. The SignalP-5.0 Server (http://www.cbs.dtu.dk/services/SignalP/) was used to predict signal peptide sequence. The Clustal X and MEGA-X programs were used for protein alignments and phylogenetic tree generation with an unrooted neighbor-joining method. The I-TASSER (http://zhanglab.ccmb.med.umich.edu/I-TASSER/) was used to predict 3D structural model.

### Transformation, pathogenicity and cell wall stress tolerance assay of *B. cinerea*

Genetic transformation of *B. cinerea* was performed as described previously (Ma *et al.,* 2017). The *bcptp1*gene deletion mutants *ΔBcPTP1-1* and *ΔBcPTP1-2*, overexpression strains OEBcPTP1-1 and OEBcPTP2, were confirmed using PCR and RT-qPCR.

Pathogenicity assays on the primary leaves of 9-days-old bean were performed as previously described (Zhu *et al.,* 2017). Conidia of indicated *B. cinerea* strains were suspended in inoculation medium (Gamborg’s B5 medium containing 2% (w/v) glucose and 10 mM KH_2_PO_4_/K_2_HPO_4_, pH6.4), the conidia were diluted to 2×10^5^ conidia/ml and leaves were inoculated with 7.5 μl of spore suspension. Plants were incubated in a humid chamber at 22°C for 72 h, and the lesion diameter was measured.

To determine the possible effect of BcPTP1 on stress tolerance and cell wall integrity, the indicated strains were inoculated on PDA plates supplemented with 0.5 mg/ml Congo Red, 0.3 mg/ml Calcofluor White, 1 M sorbitol, 1 M NaCl, 20 mM H_2_O_2_ and 0.02% SDS at 22°C, as described previously (Zhu *et al.,* 2017).

### *A. tumefaciens*-mediated transient expression and Western blotting assay

*A. tumefaciens*-mediated transient expression in *N. benthamiana* leaves was performed using agroinfiltration method as previously described (Zhu *et al.,* 2017). For extraction of fungal or plant proteins, 0.2 g of tissue was ground to powder in liquid nitrogen, suspended in 1 ml of cold lysis buffer (Beyotime Biotechnology, Shanghai, China), the samples were incubated on ice for 5 min and then centrifuged at 12,000 *g* for 10 min at 4°C, the supernatant containing soluble proteins was collected. The supernatant proteins were then mixed with 5×SDS-PAGE sample buffer (Beyotime Biotechnology, Shanghai, China), denatured by boiling for 10 min at 100°C and then separated by SDS-PAGE electrophoresis and transferred onto PVDF membranes (0.45 μm). Western blotting was analyzed using anti-GFP antibody (Beyotime Biotechnology, Shanghai, China).

### Expression and purification of recombinant BcPTP1 protein

Expression of BcPTP1 and GFP recombinant proteins were performed in *E. coli* strain Rosetta-gami (DE3) as described previously (Zhu *et al.,* 2017). Purification of recombinant proteins was performed using Glutathione Beads 4FF (Smart-Lifesciences, Changzhou, China) according to manufacturer’s instructions. The proteins were cleaned using Amicon Ultra-4 Centrifugal Filter Devices (15 ml, 10 kD; Merck Millipore) to remove the elution buffer, dissolved in phosphate-buffered saline (PBS) and stored at –80°C.

### Protein infiltration assay and induction of plant resistance by BcPTP1

To test the phytotoxic activity of BcPTP1, leaves were infiltrated with recombinant protein solution. Plants were then kept in chamber at 25°C and photographed at 10- and 15-days after treatment.

To test the of BcPTP1 on plant defense response and sensitivity to infection, *N. benthamiana* leaves were infiltrated with 10 μg/ml protein solution. The infiltrated plants were kept in a greenhouse for 48 h, and then the treated leaves were inoculated with *B. cinerea* and incubated for an additional 48 h, or used for expression analysis of defense-related genes using RT-qPCR.

### VIGS in N. benthamiana

To test whether BcPTP1-induced plant death is associated with *NbBAK1* or *NbSOBIR1* in tobacco, VIGS was used to silence the expression of *NbBAK1* or *NbSOBIR1* as described previously (Zhu *et al.,* 2017). The plasmid pTRV2 :: *GFP* was used as the control. The expression level of *NbBAK1* or *NbSOBIR1* in gene silenced *N. benthamiana* was determined by RT-qPCR analysis. Then, the *BcPTP1* was expressed using *A. tumefaciens*-mediated transient expression in the leaves of *NbBAK1* or *NbSOBIR1* silenced *N. benthamiana*. The infiltrated plants were then kept in greenhouse at 25°C and the plant death development was photographed at 10 days and 15 days after infiltration.

### Subcellular localization analysis

To analyze the subcellular localization of BcPTP1 in plant cells, the BcPTP1-GFP, BcPTP1^ΔSP^-GFP, GFP and SP^BcPTP1^-GFP sequences were cloned into binary vector pCNG between the 2×CaMV 35S promoter and NOS terminator, respectively. Transient expression was carried out using agroinfiltration. *N. benthamiana* leaves epidermal cells were harvested 3 days after agroinfiltration and plasmolyzed in 0.75 M sucrose solution, then the samples were imaged under confocal laser scanning microscope (Leica TCS SP8). Excitation wavelength of 488 nm, emission wavelength of 495-510 nm was used to detect GFP, and 650-750 nm was used for chloroplast autofluorescence.

### Statistical analysis

OriginPro 2021b (OriginLab Corporation, Northampton, MA, USA) was used for statistical tests. ANOVA (one-way, P ≤ 0.01) was used to analyze data significance. In all graphs, results were obtained from three to five independent experiments. Asterisks or different letters in the graphs indicate statistical differences at P ≤ 0.01.

## Supplemental data

**Fig. S1.**
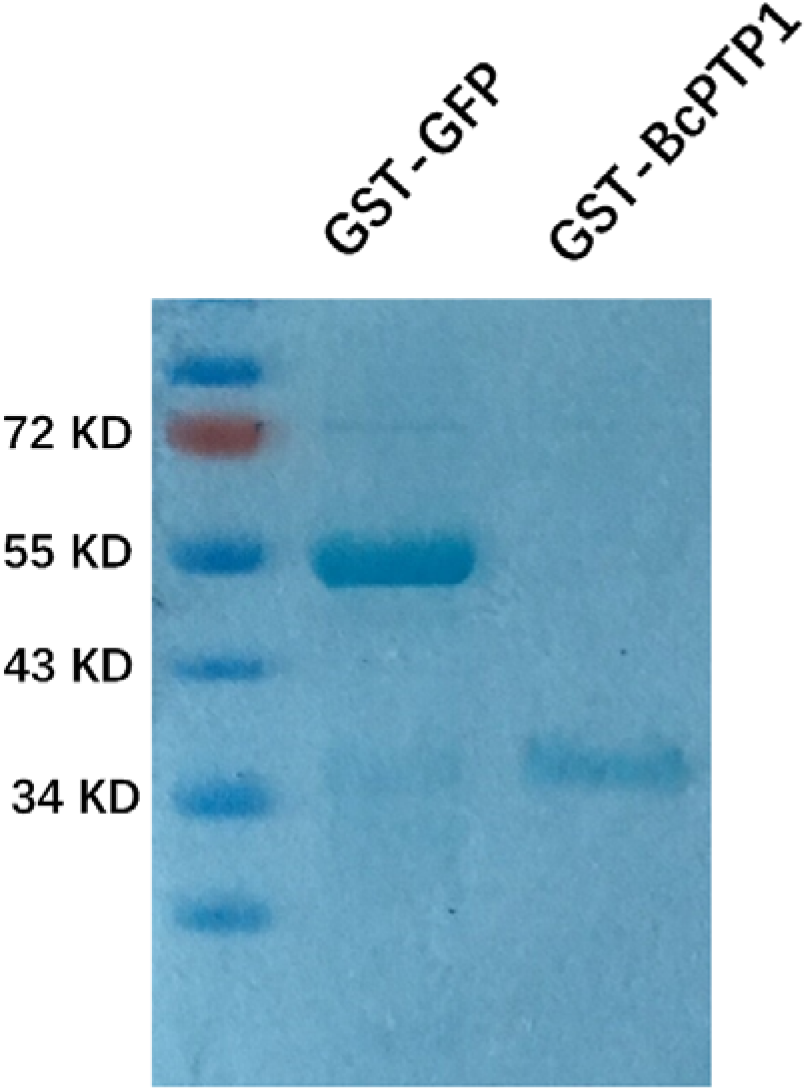
Expression of recombinant proteins. SDS-PAGE analysis of purified GST- BcPTP1 and GST-GFP proteins from *E. coli* stained with Coomassie Blue.

**Fig. S2.**
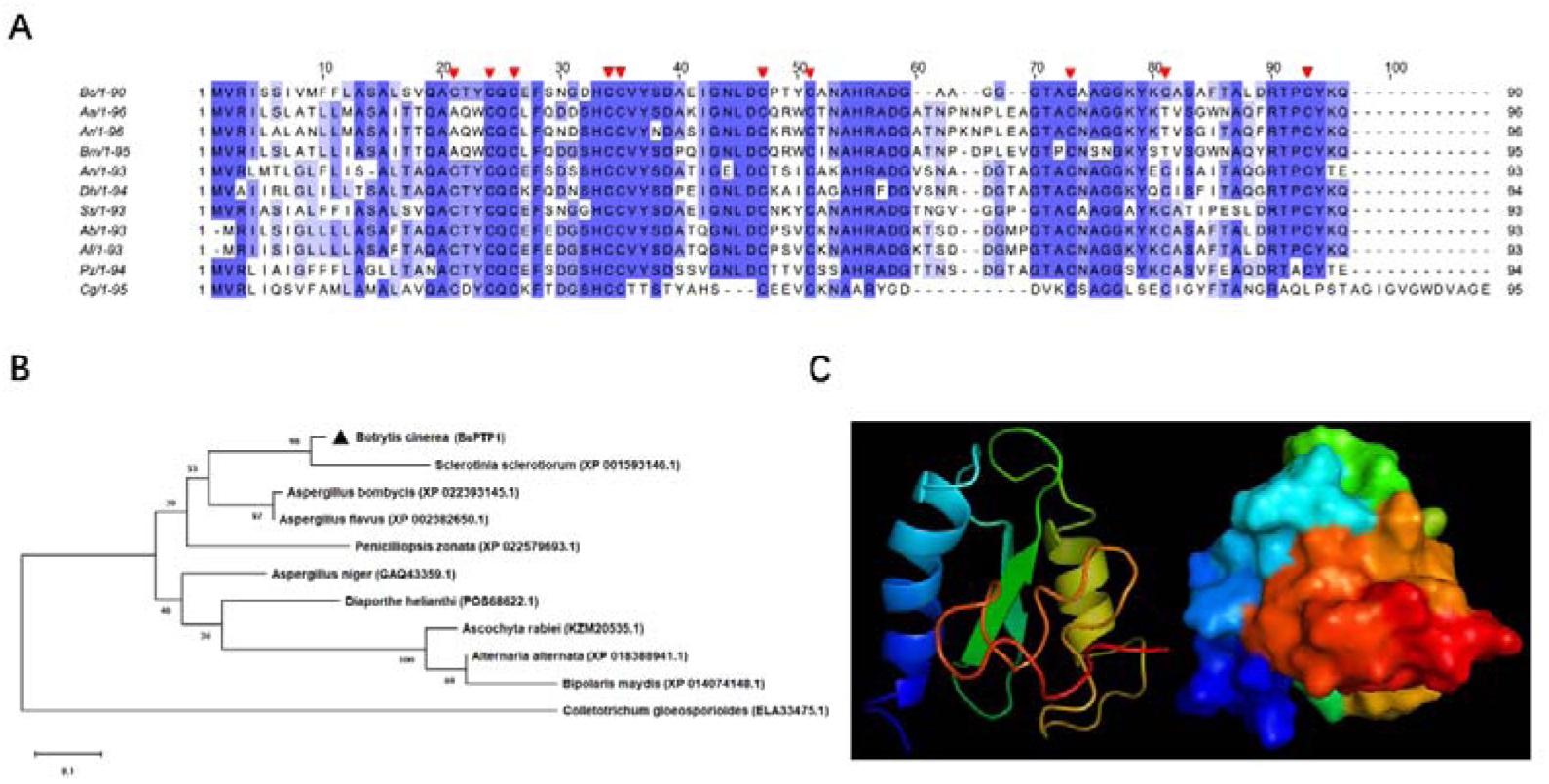
Sequence similarities between BcPTP1 and its homologs. (A) Multiple sequence alignment of BcPTP1 and its homologs. Full-length protein sequences were aligned using *Clustal W* and the alignment was edited using *Jalview*. The red triangles show the cysteine residues. Intensity of blue shading reflects the level of amino acid identity at each position. Bc: *B. cinerea* BcPTP1 (BCIN_05g03680); Ss: *S. sclerotiorum* (XP_001593146.1, E-value: 8e-45, 80% identity); Ab: *Aspergillus bombycis* (XP_022393145.1, E-value: 2e-39, 72% identity); Af: *A. flavus* (XP_002382650.1, E-value: 2e-39, 72% identity); Aa: *A. alternata* (XP_018388941.1, E-value: 1e-31, 58% identity); An: *A. niger* (G359.1, E-value: 2e-31, 61% identity); Cg: *C. gloeosporioide*s (ELA33475.1, E-value: 4e-09, 44% identity); Pz: *Penicilliopsis zonata* (XP_022579693.1, E-value: 5e-34, 63% identity); Ar: *Ascochyta rabiei* (KZM20535.1, E-value: 2e-31, 58% identity); Dh: *Diaporthe helianthi* (POS68622.1, E-value: 4e-31, 61% identity); Bm: *Bipolaris maydis* (XP_014074148.1, E-value: 2e-29, 55% identity). (B) Phylogenetic analysis of BcPTP1 and its homologs from other fungi. The full-length protein sequences were analyzed using *MEGA X* with Unrooted neighbor-joining bootstrap (1,000 replicates). The black triangle mark location of BcPTP1. A scale bar at the lower left corresponds to a genetic distance of 0.1. (C) 3D structural models of BcPTP1, predicted using *I-TASSER* and further analyzed by *PyMOL* software. Left: cartoon models; right: surface models.

**Fig. S3.**
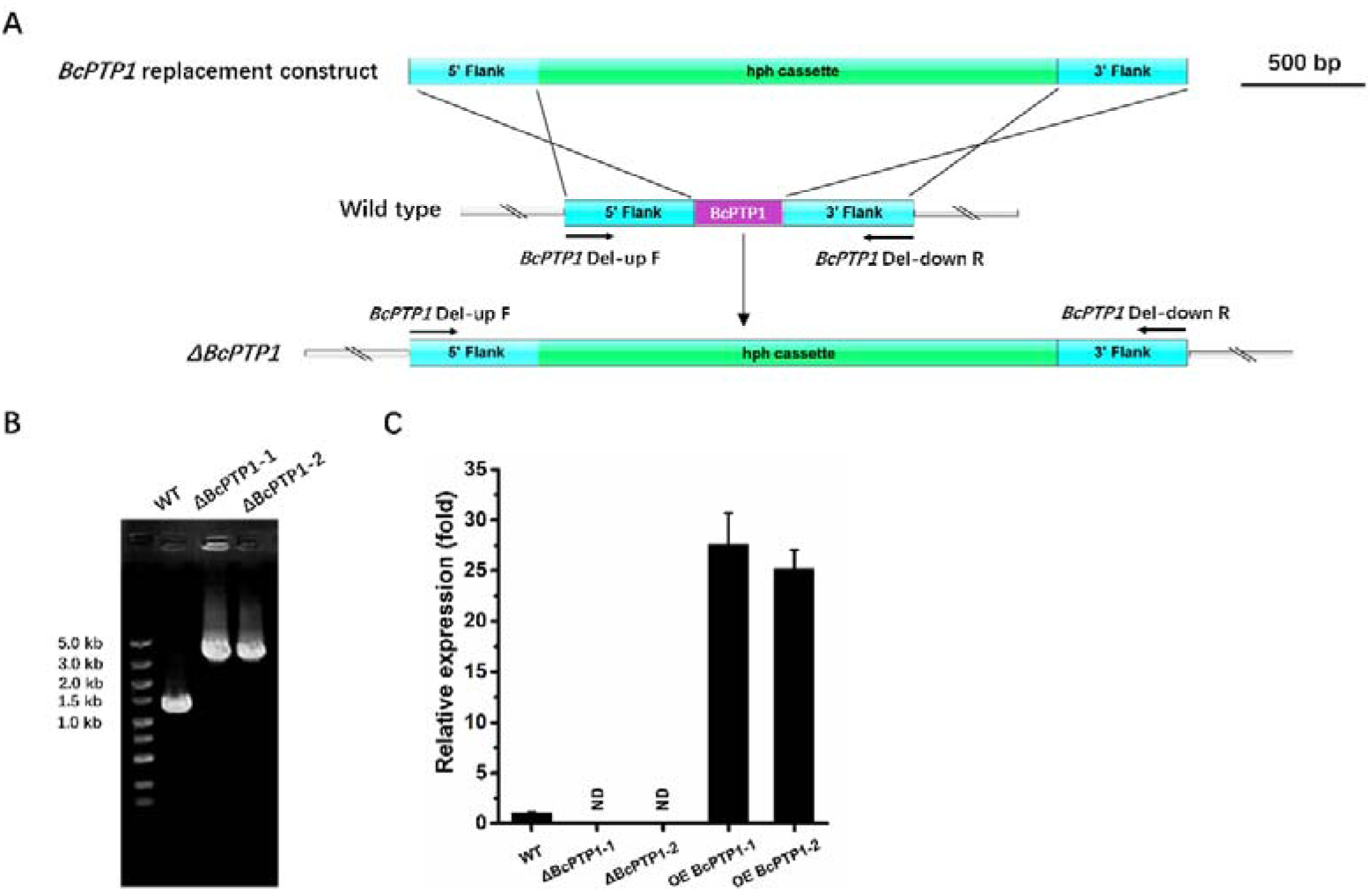
The *bcptp1* gene deletion strategy and confirmation in *B. cinerea*. (A) Strategy used to generate the *bcptp1* gene deletion mutants. The deletion construct used to transform the wild type strain contained the hygromycin resistance (*hph*) cassette flanked by upstream and downstream sequence of the *bcptp1* gene. The positions of the PCR primers used to verify the deletion transformants are indicated (*BcPTP1* Del-up F and *BcPTP1* Del-down R). The scale bar indicates 500 bp. (B) PCR amplifications carried out to verify the *bcptp1* deletion mutants using the 5’ flank For and 3’ flank Rev primers. As templates, genomic DNA from either the wild type strain or the *bcptp1* deletion mutants Δ*BcPTP1*-1 and Δ*BcPTP1*-2, was used as indicated. (C) RT-qPCR carried out to analyze the expression levels of the *bcptp1* gene in the wild type strain, the *bcptp1* deletion mutants Δ*BcPTP1*-1 and Δ the *bcptp1* overexpression strains OE *BcPTP1*-1 and OE *BcPTP1*-2. The strains cultured on PDA at 22°C for 72 h were used for RT-qPCR analysis. The relative levels of transcript were calculated using the comparative Ct method. The *bcptp1* gene expression level in wild type strain was set as level 1. The transcript level of the *B. cinerea bcgpdh* gene was used to normalize different samples. Data represent means and standard deviations of three independent replicates. ND=not detected.

**Fig. S4.**
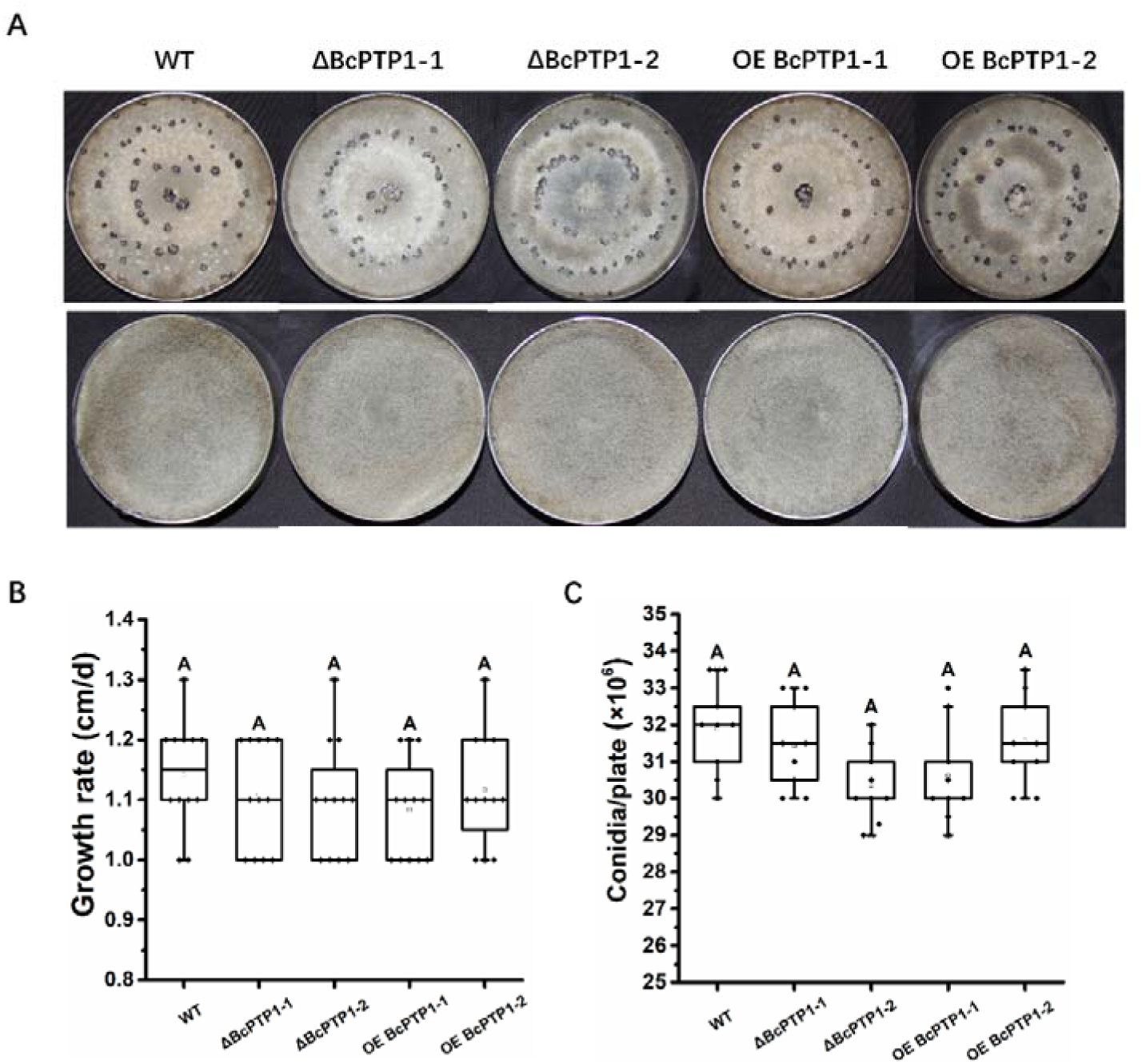
Phenotypes of *B. cinerea* wild type, and the *bcptp1* deletion and overexpression strains. (A) Colony morphology, sporulation and sclerotia formation. Top: colonies on PDA at 22°C for 15 days in complete darkness. Bottom: *B. cinerea* strains on PDA plates at 22°C for 7 days with continuous fluorescent light. (B) Hyphal growth rate. *B. cinerea* strains were grown on PDA plates at 22°C with continuous fluorescent light. Radial growth was measured every day for 5 days and growth rate was calculated. Data were obtained from three independent experiments, with four replicates in each experiment. In box plots, whiskers indicate the minimum and maximum values, the line indicates the median, the box boundaries indicate the upper (25th percentile) and lower (75th percentile) quartiles, all data are plotted as black dots. Same letters in the graph indicate no statistical differences at P ≤ 0.01 using one-way ANOVA one-way. (C) Conidial production of indicated strains cultured on PDA plates at 22 °C for 7 days. Conidiation of each strain was determined by collecting and counting conidia with a hematocytometer. Data were obtained from three independent experiments, with three replicates in each experiment. In box plots, whiskers indicate the minimum and maximum values, the line indicates the median, the box boundaries indicate the upper (25th percentile) and lower (75th percentile) quartiles, all data are plotted as black dots. Same letters in the graph indicate no statistical differences at P ≤ 0.01 using one-way ANOVA.

**Fig. S5.**
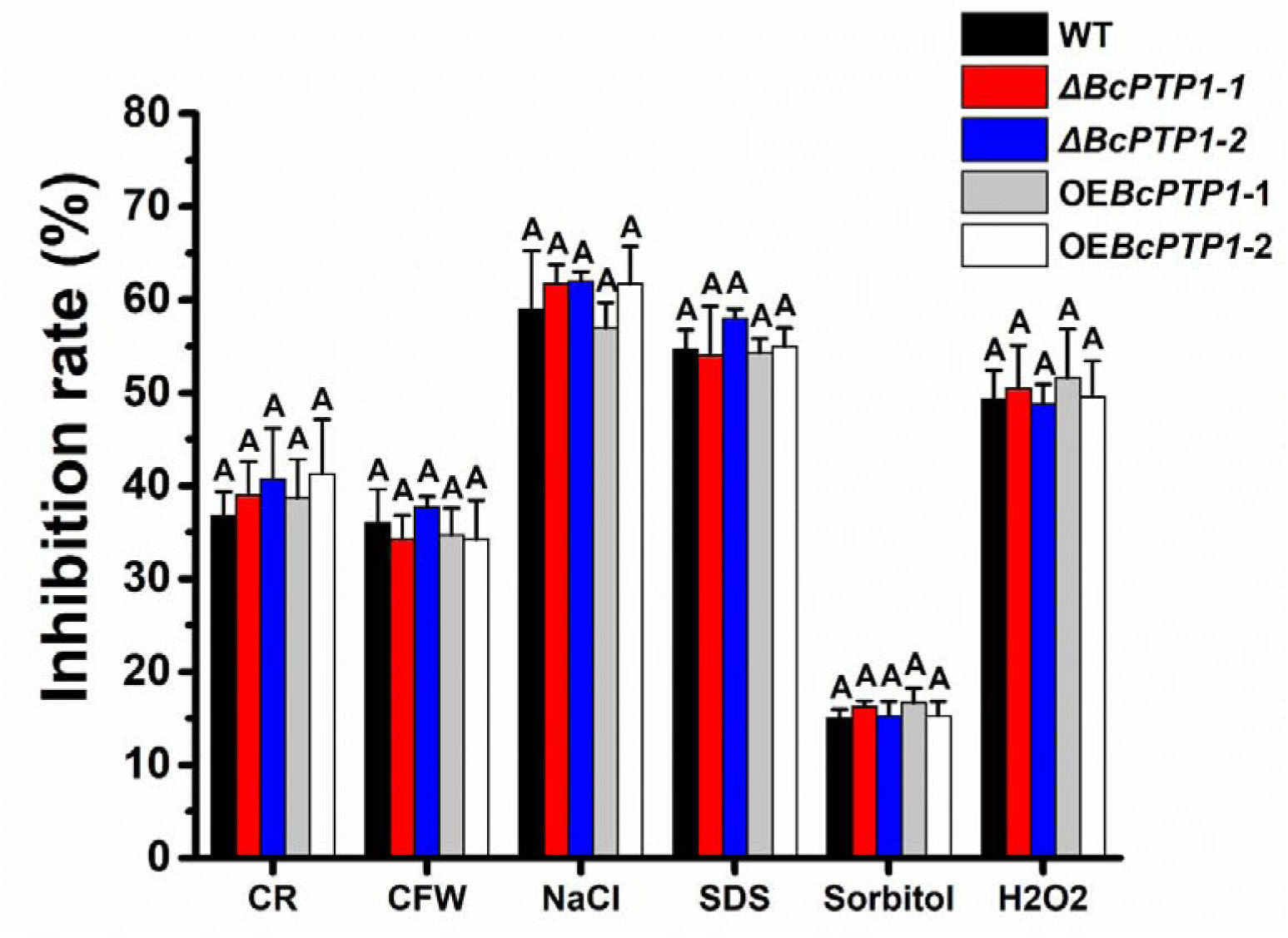
*bcptp1* deletion does not affect stress tolerance of *B. cinerea*. Inhibition rate of the radial growth of wild type strain, *bcptp1* deletion mutants and *bcptp1* overexpression strains were analyzed on PDA plates supplemented with 0.5 mg/ml Congo Red, 0.3 mg/ml Calcofluor White, 1 M sorbitol, 1 M NaCl, 20 mM H_2_O_2_ and 0.02% SDS at 22°C, respectively. Data represent means and standard deviations from three independent experiments, each with three replications. Same letters in the graph indicate no statistical differences at P ≤ 0.01 using ANOVA (one-way).

**Fig. S6.**
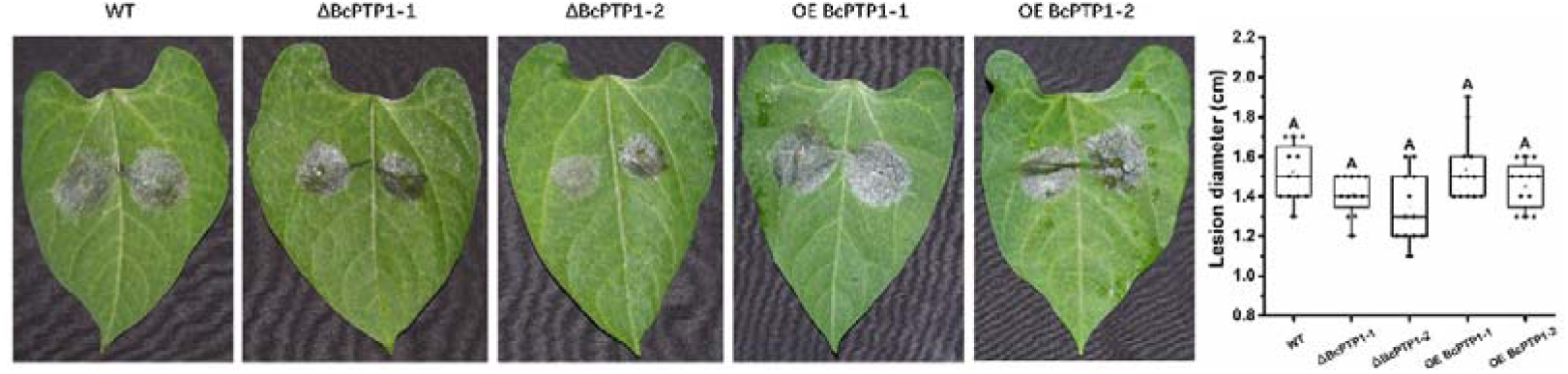
Pathogenicity analysis of *B. cinerea* strains. Beans leaves were inoculated with 7.5 μl of conidia suspension (2×10^5^ conidia/ml). The plants were incubated in a humid chamber at 22°C for 72 h, photographed and lesion size was determined. Data were obtained from three independent experiments, with four replicates in each experiment. In box plots, whiskers indicate the minimum and maximum values, the line indicates the median, the box boundaries indicate the upper (25th percentile) and lower (75th percentile) quartiles, all data are plotted as black dots. Same letters in the graph indicate no statistical differences at P ≤ 0.01 using ANOVA (one-way).

**Table S1.**
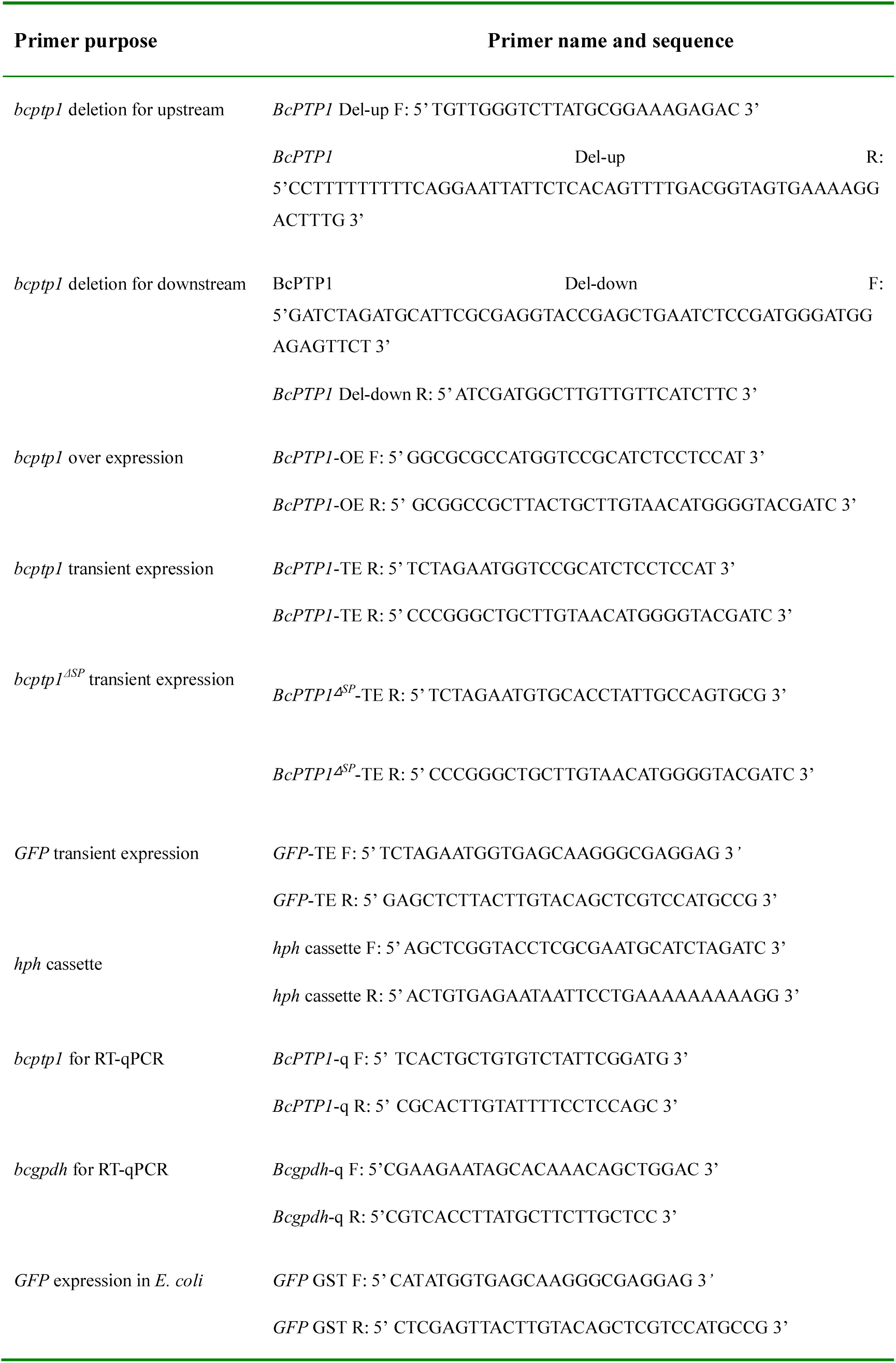
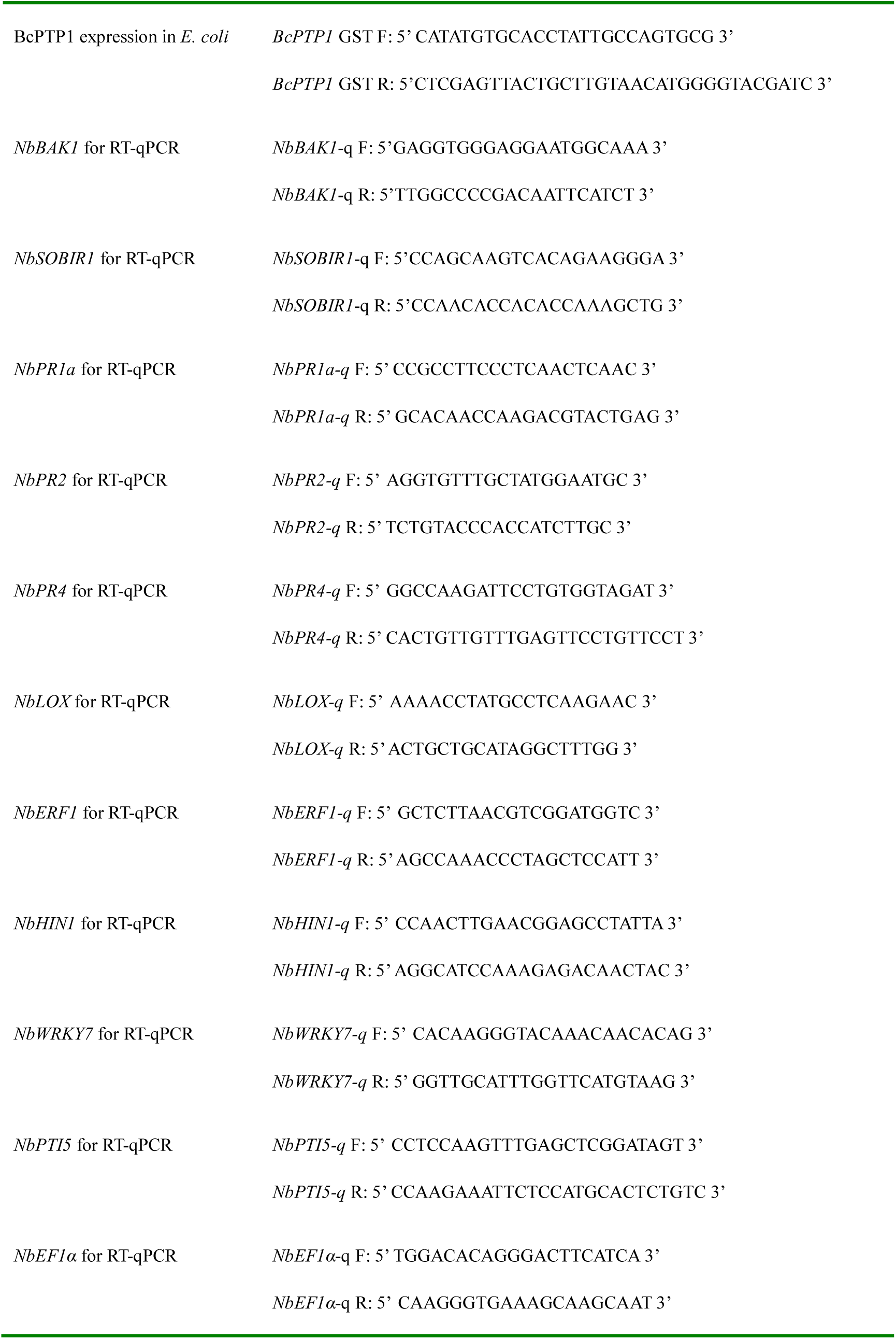
Primers used for vector construction and PCR.

## Notes

### Competing Interest Statement

The authors have declared no competing interest.

